# Working Memory of Multi-Object Scenes in Primate Frontal Cortex

**DOI:** 10.64898/2026.01.27.702062

**Authors:** Nicholas Watters, Jack Gabel, Joshua Tenenbaum, Mehrdad Jazayeri

## Abstract

Working memory allows primates to reason about complex scenes, yet how the brain maintains multiple objects in memory simultaneously remains unclear. Competing theories propose that objects are stored in discrete slots^1,2^, represented dynamically through switching^3–6^, or encoded by weighted combinations of single-object representations^7–11^. We formalized these hypotheses in terms of their quantitative predictions at the level of single neurons and tested them against densely recorded neural data from the dorsomedial frontal cortex and frontal eye field of monkeys trained to perform a novel multi-object working-memory task. Across cross-validated neural data, a Gain model, where population activity reflects weighted compositions of individual object responses, consistently outperformed Slot and Switching models. Trial-specific gain estimates predicted behavioral errors and reaction times, indicating that these latent weights capture meaningful fluctuations in memory fidelity. All results replicated in an independent dataset with different spatial configurations. Together, our work provides a rigorous framework to adjudicate a longstanding debate about how the frontal cortex retains multiple objects, identifying a weighted-sum representation as the format that best explains the neural data.

## Introduction

Primates routinely reason about complex visual scenes that contain multiple objects, each with distinct identities, locations, and behavioral relevance. This capacity depends critically on a memory system that can hold information about multiple objects. Yet despite decades of research, how the brain represents multiple objects simultaneously remains unresolved.

A central challenge is that multi-object working memory is inherently dynamic and variable. The fidelity with which individual objects are retained can fluctuate across time and trials, even when task demands are identical. These fluctuations are largely covert, distributed across large neural populations, and only indirectly reflected in behavior. As a result, competing theories of multi-object working memory have been difficult to adjudicate.

Broadly, three classes of hypotheses have dominated the literature. The Slot hypothesis proposes that working memory consists of a limited number of discrete representational slots, or “object files,” each capable of storing a single object independently of the others^1,2,12–15^. The Switching hypothesis proposes that neural activity represents only one object at a time, rapidly alternating between objects through temporal multiplexing^3–5,16–20^. Finally, the Gain hypothesis posits that multiple objects are represented simultaneously through a weighted combination of single-object neural responses, with the weights reflecting a shared, distributed resource that modulates the fidelity of each object’s representation^7–11,21–24^.

Although these hypotheses make qualitatively different claims about representational format, they share a critical feature: each relies on latent variables that can vary from trial to trial even when the memory demands are the same. In the Slot hypothesis, these variables determine how objects are assigned to slots; in the Switching hypothesis, they specify which object is represented at each moment in time; and in the Gain hypothesis, they correspond to object-specific weights.

These covert latent variables are difficult to infer from behavior alone: Behavior measurements provide only an indirect readout of internal states, collapsing over trial-to-trial variability in the underlying representations. Consequently, these hypotheses all can give rise to similar behavioral error patterns, and testing them using behavior alone has been a source of debate for decades^25–32^.

Testing these hypotheses at the neural level has also proven challenging. Latent variables that vary on a trial-by-trial basis are difficult to infer reliably from noisy spiking activity, particularly when recordings sample only small numbers of neurons. Consequently, neural evidence has been interpreted in support of each hypothesis, often without a common quantitative framework for comparison^33–38^.

Here, we address this longstanding impasse using a combination of behavioral, technical, and conceptual innovations. We trained monkeys to perform a novel multi-object working memory task and recorded the simultaneous activity of thousands of neurons in dorsomedial frontal cortex (DMFC) and frontal eye field (FEF), two regions implicated in visual working memory, attention, and saccade planning^39–47^. We then formalized the Slot, Switching, and Gain hypotheses as explicit encoding models that predict single-neuron firing rates during the memory delay, while allowing for time-dependent and trial-specific latent variables.

This integrative approach enables direct, rigorous, and head-to-head comparison of competing theories. We find that a Gain model, in which population activity reflects trial-specific weighted combinations of individual object representations, consistently outperforms Slot and Switching models. Moreover, the inferred gain weights predict trial-by-trial variability in behavioral accuracy and reaction time, linking latent neural representations to memory fidelity and decision speed. These results replicate across animals and across an independent dataset with different stimulus configurations.

Together, our findings indicate that multi-object working memory in primate frontal cortex is best described by a gain-modulated compositional code, rather than by discrete slots or rapid serial switching. More broadly, our work demonstrates how formalizing cognitive hypotheses within a shared computational framework, combined with large-scale population recordings, can reveal the representational format of latent variables even when those variables are subject to covert fluctuations.

## Results

### Task, Behavior, and Hypotheses

We trained two rhesus macaques on a multi-object working memory task (Fig. 1a). Each trial began with central fixation, followed by a 1-second presentation of 1–3 visual stimulus objects (fruit images) at random locations. The objects then disappeared for a 1-second delay, after which one of the objects reappeared at fixation as a cue. The monkey had to then make a saccade to the location of the matching stimulus object to receive a juice reward. The stimulus object positions and identities were independently randomized across trials, and the cue’s identity was randomly selected on each trial from the set of stimulus object identities on that trial. Thus, correct performance required maintaining both the identities and positions of multiple objects during the delay. Animals learned the task and exhibited variable reaction times, with error rates increasing as the number of objects grew (Fig. 1b–c; Extended Data Fig. 1). However, performance remained far above chance for two- and three-object stimuli (Fig. 1b), indicating that both monkeys could maintain multiple objects in working memory. Accordingly, we proceeded with electrophysiology recordings in the frontal cortex to test the Slot, Switching, and Gain hypotheses (Fig. 1d).

**Figure 1:**
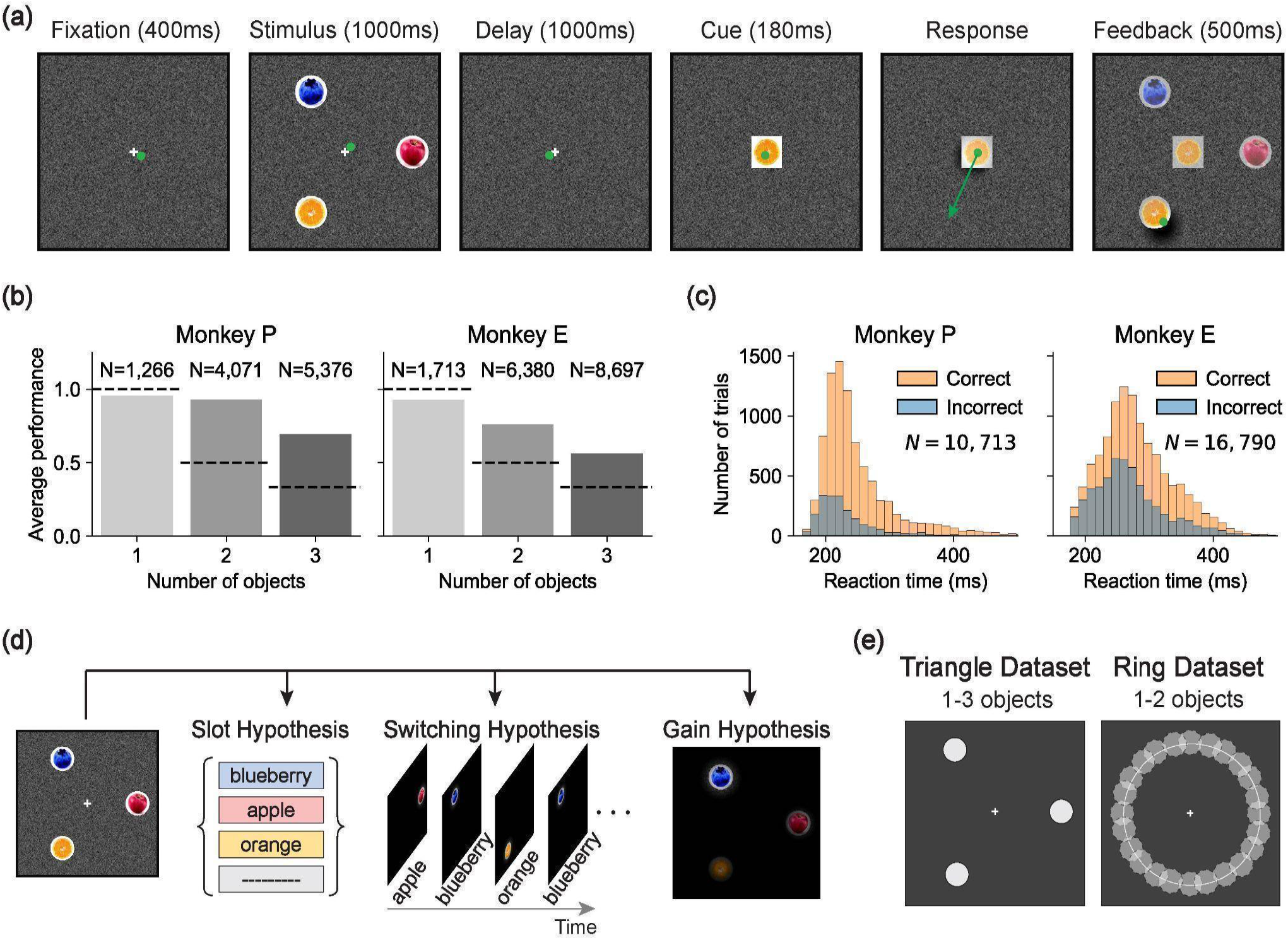
Task, behavior, and hypotheses. **(a)** Example trial (left to right). In all panels, the green dot shows the monkey’s gaze position and is not displayed during the task. Gaze fixation near the display center must be held from trial onset through cue presentation. The green arrow in the response phase depicts a saccade vector and is not displayed during the task. After the response saccade, all stimulus objects are revealed with partial opacity for visual feedback and a reward is delivered if the response was correct (near the stimulus object matching the cue). **(b)** Performance for Monkey P (left) and Monkey E (right), measured as the average fraction of correct responses for all physiology sessions, conditioned on the number of objects in the stimulus (x-axis). Each dashed line shows expected performance if the monkey responded by saccading to the location of a random stimulus object **(c)** Reaction time histograms for Monkey P (left) and Monkey E (right) over all physiology sessions. Reaction time is measured from cue onset to onset of response saccade. Trials in which a saccade was made less than 180ms after cue onset were considered as broken fixation and are not shown. **(d)** Example stimulus (left) and schematics of each hypothesis (right) for working memory of this stimulus. In the Slot hypothesis, a set of four object slots is shown, three of which are occupied by stimulus objects. In the Switching hypothesis, only one object is active in memory at any moment in time, but this active object changes through time. In the Gain hypothesis, working memory resources are distributed across objects but not necessarily equally, leading to some objects being retained with high fidelity and others with low fidelity. **(e)** For the Triangle dataset (left), each stimulus had 1-3 objects with positions drawn from the vertices of an equilateral triangle (positions of the light circles). For the Ring dataset (right), each stimulus had 1-2 objects with positions drawn from a uniform distribution on a circle. The results in panels (b) and (c) include data from both the Triangle and Ring datasets.

### Frontal Cortex Neural Activity

We used high-density probes (Neuropixels) to record spiking activity simultaneously in the dorsomedial frontal cortex (DMFC) and frontal eye field (FEF) while monkeys performed the task. We focused on these areas as they are thought to support visual working memory function, attention, and saccade planning^39–46^.

We collected two datasets (Fig. 1e): a **Triangle dataset**, with object positions restricted to three spatial locations, and a **Ring dataset**, with positions sampled uniformly from a circle. We treated the two datasets independently: all analyses and model comparisons were first performed on the Triangle dataset, and the same models and procedures were subsequently applied to the Ring dataset to assess validity.

### The Triangle Dataset

We recorded 4,688 well-isolated neurons (2,937 from DMFC and 1,751 from FEF) in the Triangle dataset across 24 sessions (see Methods). In both brain areas, neurons were modulated during both the stimulus and delay phases of the trial and exhibited diverse temporal dynamics (Fig. 2a). We found that neurons were typically active in both stimulus and delay phases (Extended Data Fig. 2). We also found that neural activity contained information about object positions and identities (Extended Data Fig. 3a-b). A Receiver Operating Characteristic analysis of firing rates in the delay phase revealed a wide range of selectivities for position and identity across neurons (Fig. 2b).

**Figure 2:**
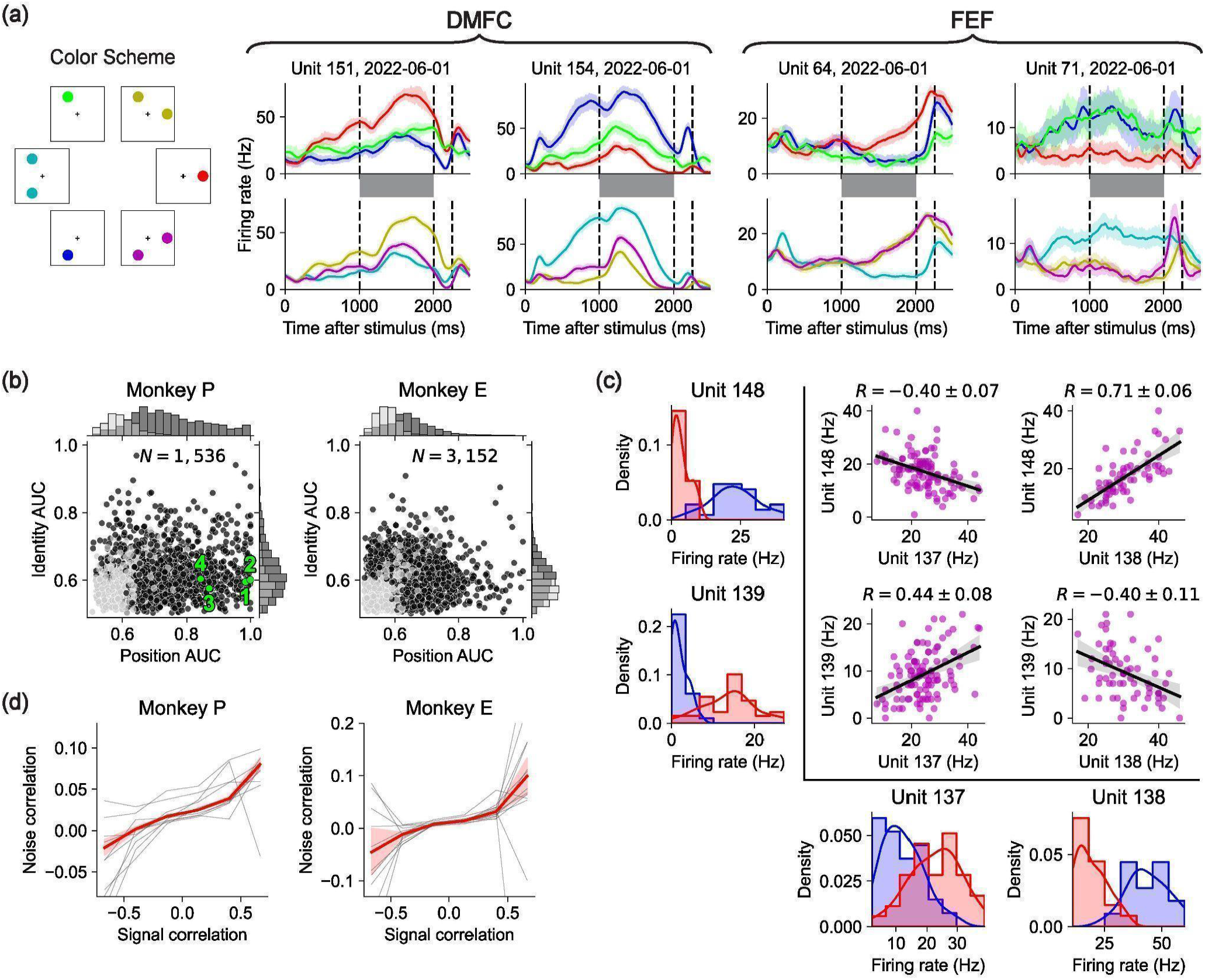
Neural data in the Triangle dataset. **(a)** Firing rate of 4 example neurons during the trial for 1-object (top) and 2-object (bottom) conditions in the Triangle dataset, marginalizing over stimulus object identities. Lines show mean firing rates smoothed by convolution with a triangular kernel with 200ms support, and errorbands show 95% confidence interval of the mean. Vertical dashed lines show time of delay onset, cue, and mean response time. Gray shaded regions highlight the delay phase. The left panel shows the color scheme for different conditions. **(b)** Position- and identity-selectivity of every recorded unit using area under the receiver operator characteristic (AUC) for 1-object trials. For position AUC we marginalized data over identity, and vice versa. AUC values are cross-validated (dark/light gray: significant/insignificant at p-value of 0.05). Green values correspond to the units shown in panel (a), indexed left-to-right. **(c)** Firing rate fluctuations and trial-by-trial co-fluctuations of four simultaneously recorded units. The histograms on the left and bottom show the delay-phase firing rate distribution for the four units in two specific 1-object configurations (see color scheme in panel a). The smooth lines in the histograms are kernel density estimates of the continuous distributions. The scatterplots show the delay-phase firing rate of one neuron with respect to another on a trial-by-trial basis. Each magenta point corresponds to one trial when both objects were present. The black lines are regression lines and the shaded region shows the 95% confidence interval of the regression. All plots in this panel marginalize over identity, yet all neurons in this panel do not have significant tuning to identity (p > 0.05) and these results hold when conditionalized on object identity (Extended Data Fig. 4). **(d)** The relationship between noise and signal correlations for all pairs of simultaneously recorded neurons in the Triangle dataset. Given a pair of neurons, signal correlation is measured as the correlation coefficient between their mean delay-phase firing rates across all 1-object trials, and noise correlation is measured as the mean over each 2-object stimulus (conditionalizing on identity) of the correlation coefficient of their delay-phase firing rates. Gray lines are means per session. Red lines show grand means over all pairs of units on all sessions, with 95% confidence interval of the mean shaded.

Despite this heterogeneity in single-neuron selectivities and modulation, we observed a systematic effect that bears on compositional representations of multiple objects during the delay phase: Pairs of neurons with similar selectivities on 1-object trials tended to have positive noise correlations on 2-object trials — for example, if two neurons preferred object A at position 1 over object B at position 2, they tended to have positive noise correlations for the composite stimuli consisting of both A at position 1 and B at position 2. Conversely, dissimilarly tuned pairs of neurons tended to have negative noise correlations. This pattern was evident both at the level of individual neuron pairs (Fig. 2c) and in aggregate across the population (Fig. 2d).

This pattern of noise correlations is consistent with an interpretation based on trial-by-trial fluctuations in covert attention. Suppose neurons have receptive fields defined jointly over stimulus position and identity. For any given 2-object stimulus, neural activity may be more strongly driven by one object on some trials and by the other on different trials, depending on the allocation of attention to objects. This variability would lead neuron pairs with similar selectivity (i.e., positive signal correlations) to exhibit positive noise correlations, and those with opposing selectivity (negative signal correlations) to exhibit negative noise correlations, as we observe. While this attentional interpretation is plausible and compatible with prior studies of attention and normalization in neural activity^7,8,46,48^, it is not diagnostic of any one hypothesis. In the Slot hypothesis, this could arise from variability in the assignment of objects to memory slots across trials. In the Switching hypothesis, it could arise from fluctuations in the time devoted to each object within a trial. In the Gain hypothesis, it could arise from trial-by-trial variations of the gain factors applied to each object representation. Therefore, while this effect underscores the importance of accounting for trial-level neural variability, distinguishing among the hypotheses requires a formal computational modeling approach.

### Formalizing Hypotheses

To test which of the three hypotheses — Slot, Switching, or Gain — best explains how the frontal cortex stores multiple objects in working memory, we formalized each hypothesis as a distinct encoding model (Fig. 3). Each model predicts the firing rate of every neuron *n*, on every trial *k*, at every time point *t* during the delay phase, according to the assumptions of the corresponding hypothesis.

**Figure 3:**
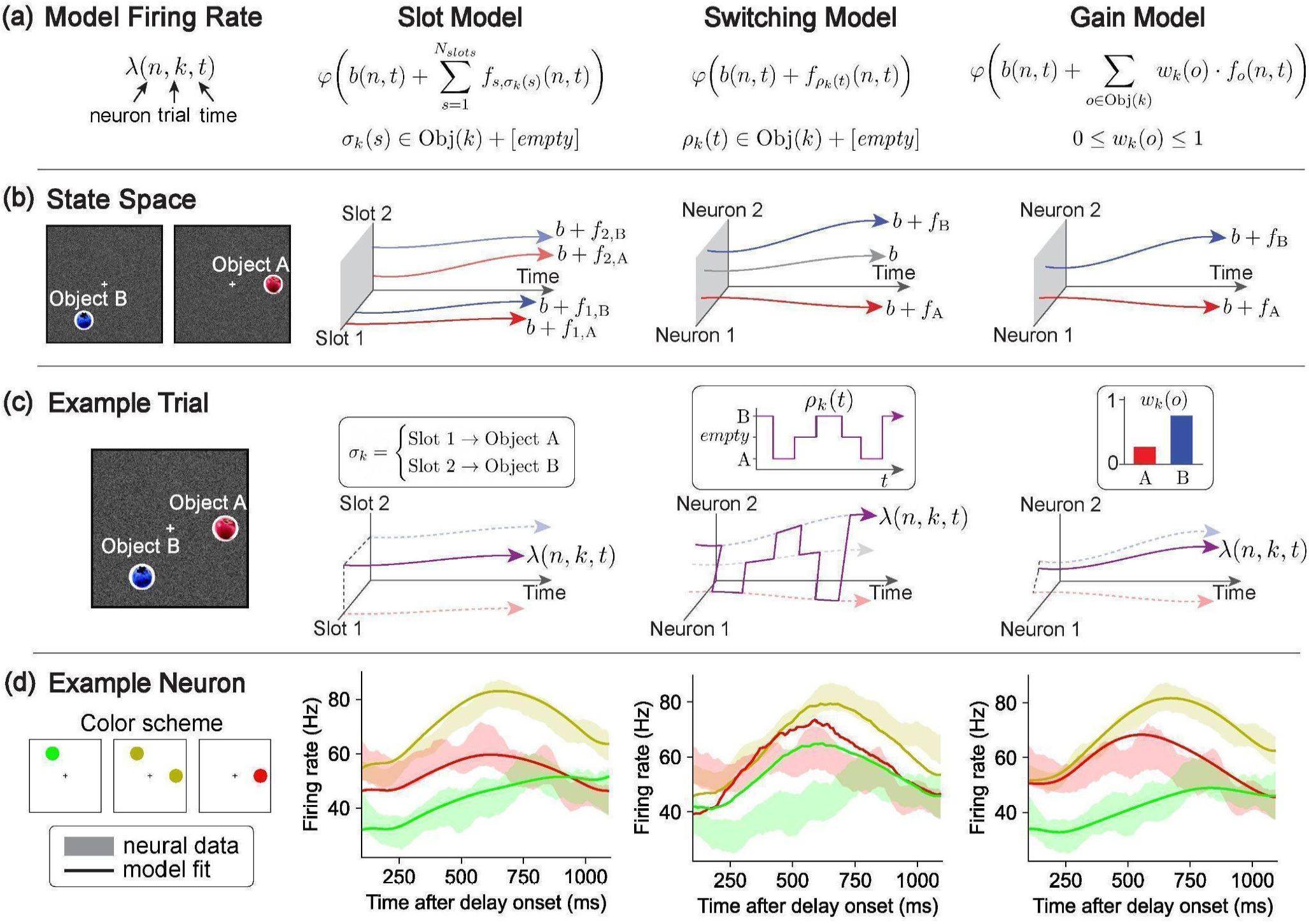
Hypothesis formalizations. **(a)** Each model parametrizes the firing rate of each neuron at every timepoint in every trial. Different models have different constraints imposed by their formulation. **(b)** (far left) Two example 1-object stimuli. (middle left) Schematic neural state space structure for the Slot model for the two 1-object stimuli. Each slot is an independent axis in neural state space. Each slot has embeddings for each object, which may vary in time. (middle and far right) Schematic neural state space structure for the Switching and Gain models for the same two 1-object stimuli. **(c)** (far left) Composite 2-object stimulus composed of the objects in panel (b). (middle left) Example schematic neural activity of the Slot model for the composite stimulus. One object is allocated to each slot. The total neural activity (magenta) is the sum of the activities in the two slots (dashed arrows). Inset shows the allocation mapping. (middle right) Example schematic neural activity of the Switching model. Neural activity (magenta) switches through time between the embedding for each object (blue and red dashed arrows) and baseline activity (gray dashed arrow). Inset shows the time-varying indicator variable determining the active object. (far right) Example schematic neural activity of the Gain model. Neural activity (magenta) is the weighted sum of the responses to each individual object (blue and red dashed arrows). Inset shows the gain coefficients. **(d)** PSTHs for an example neuron. (far left) Color scheme, as in Fig. 2a. (middle left) Shaded bands show 95% confidence intervals for each condition in the color scheme of mean delay-phase firing rate of an example unit (Monkey P, 2022-06-01, Unit 141). Lines show the mean delay-phase firing rate for each condition of the corresponding unit predicted by the Slot model. (middle right and far right) Analogous plots for the Switching model and Gain model.

#### Slot hypothesis

Under the Slot hypothesis, working memory has *N*_*slots*_ discrete slots and each object is encoded by activity in one slot. Prior implementations of the slot hypothesis define each slot as a disjoint set of neurons^49–51^. We do the same, though we found that relaxing this assumption and allowing the slots to be linear subspaces in neural activity space does not qualitatively change our results (Extended Data Fig. 5). The core of the Slot hypothesis is an assignment of slots to objects, which may vary across trials. Let σ_*k*_ (*s*)∈*Objects*(*k*) + [*empty*] denote this assignment on trial *k*, where *s*∈[1, …, *N*_*slots*_] is a slot, *Objects*(*k*) is the set of stimulus objects on trial *k*, and an *empty* allocation means that the slot does not encode any object (which may happen for trials with fewer objects than *N*_*slots*_).

We then let the firing rate of each neuron *n* at time *t* during the delay phase be the sum of an object-independent baseline *b*(*n, t*) and the activity driven by the content of the slot to which neuron *n* belongs. Each slot has the ability to encode any object, so for each object *o* and slot *s* there is a time-varying embedding function *f*_*s,o*_ (*n, t*) that expresses the activity of each neuron *n* due to the memory of object *o* in slot *s*. Because each slot is a disjoint set of neurons, *f*_*s,o*_ (*n, t*) = 0 for all neurons *n* that do not belong to slot *s*. Finally, because neurons cannot have negative firing rates, we apply a softplus nonlinearity to all model neurons 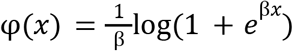 for β = 5. With these considerations, the Slot model is formulated as follows (Fig. 3a):

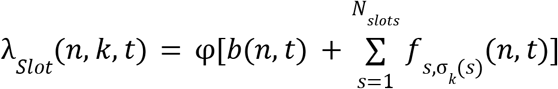

Where λ_*Slot*_ (*n, k, t*) is the model-predicted firing rate of neuron *n* at time *t* in the delay phase of trial *k*.

#### Switching hypothesis

Under the Switching hypothesis, neural activity encodes at most one object (i.e., “active object”) at any point in time, but the active object may change through time during the delay phase. The core of this hypothesis is a switching function ρ_*k*_ (*t*)∈*Objects*(*k*) + [*empty*], which determines which object, if any, is represented at any point in time *t* for each trial *k*. We then let each neuron’s firing rate be the sum of a baseline *b*(*n, t*) and the embedding of the active object. We let *f*_*o*_ (*n, t*) denote the embedding of object *o* and *f*_*empty*_ (*n, t*) denote modulation when no object is active. Accordingly, the Switching model is formulated as follows (Fig. 3a):

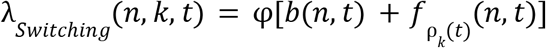

#### Gain hypothesis

Under the Gain hypothesis, neural activity for multiple objects in working memory is driven by a weighted sum of the activity associated with each individual object.

The core of this hypothesis is a gain factor on each object on each trial *w*_*k*_ (*o*)∈[0, 1], which captures the relative influence of each object on neural activity. We then let each neuron’s firing rate be the sum of a baseline *b*(*n, t*) and the weighted sum of the embedding of each object. Letting *f*_*o*_ (*n, t*) denote the embedding of object *o*, the Gain model is formulated as follows (Fig. 3a):

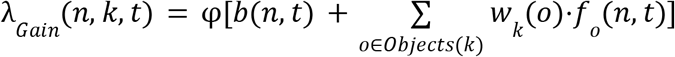

### Modeling Results

We fit each model to our neural spiking data using maximum Poisson likelihood and compared model likelihoods based on held-out neural data (see Methods for details). Across sessions, the Gain model had a higher cross-validated likelihood than the Slot and Switching models (Fig. 4a). This held for 88% of sessions, including all sessions with at least 100 simultaneously recorded units. The results were consistent across both DMFC and FEF (Extended Data Fig. 6).

**Figure 4:**
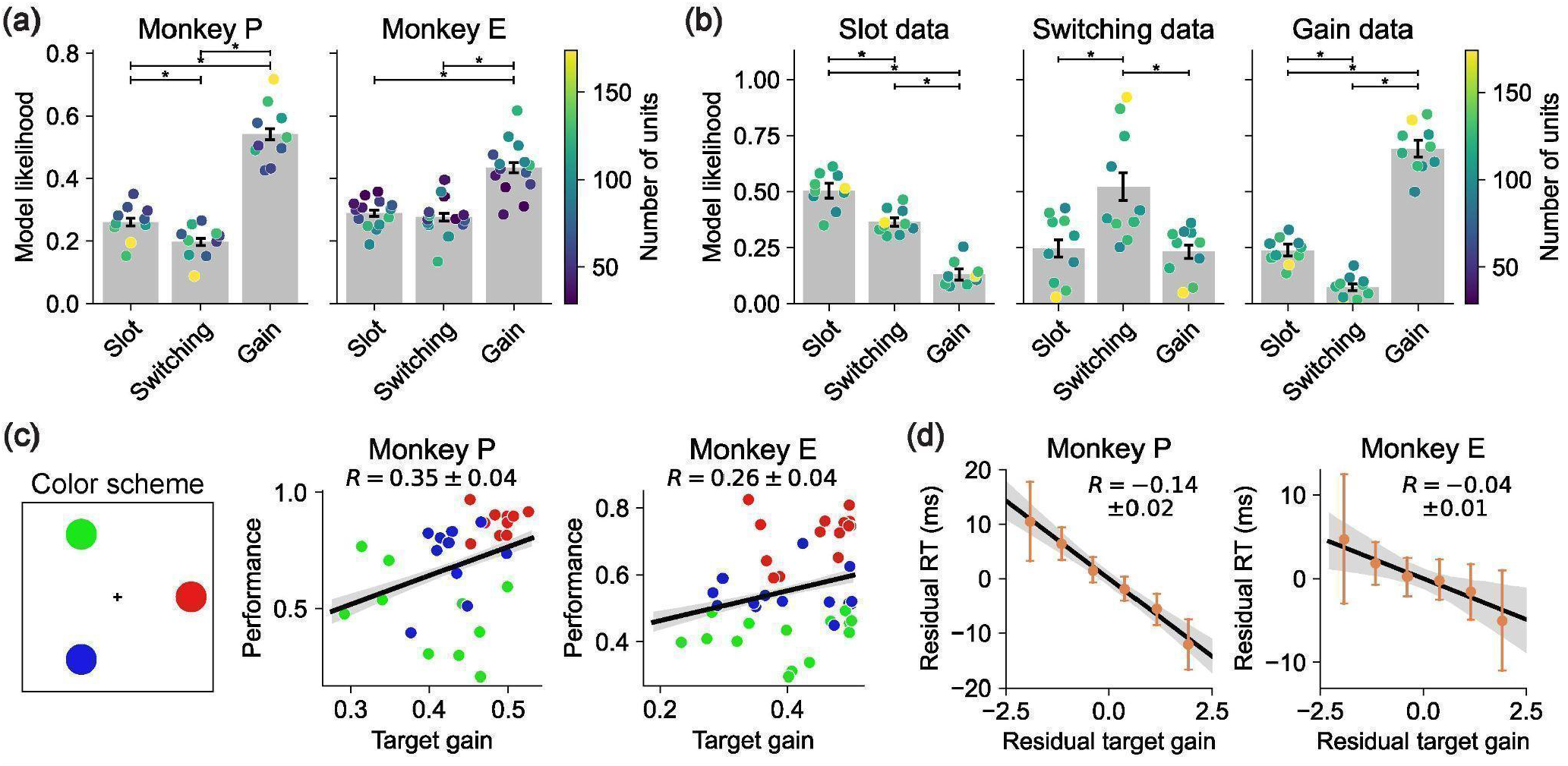
Modeling results. **(a)** Model likelihood for the Slot, Switching, and Gain models. Bar height show mean cross-validated bootstrapped model likelihoods, with errorbars showing 95% confidence interval of these means. Each scatterplot point shows the model likelihood for one session, averaged over cross-validated bootstraps and colored according to the number of simultaneously recorded units in that session. Significance brackets above show statistically significant (p < 0.05) difference of the mean. **(b)** Model identifiability. Each plot is the same style as panel (a) but uses synthetic data generated from one of the three models after fitting to neural data, and restricted to only ten sessions of data (the five sessions with the most neurons from each monkey). **(c)** *(left)* Color scheme for the three object locations in the Triangle dataset. *(middle and right)* Neurally-inferred gain values of the target object (x-axis) versus performance (y-axis) for 3-object trials. Each point represents one target location determined by the hue (left panel) for one session, averaged over random seeds. Regression line is best-fit linear regression to the mean performance per session, target location, and random seed, with shaded 95% confidence interval. The regression is significantly positive for both monkeys ( *R* = 0. 35±0. 04, *p* < 10^−9^ for Monkey P; *R* = 0. 26±0. 04, *p* < 10^−7^ for Monkey E). **(d)** Reaction time is inversely correlated with the gain of the target object for correct 3-object trials. To compute the normalized residual target gain (x-axis) for a trial, we normalize the target object’s gain (averaged over random seeds) such that the distribution of all target object gains for trials with the same stimulus, cue, and session have mean zero and variance 1. The residual reaction time for a trial is similarly de-meaned conditioned on stimulus, cue, and session, but is un-scaled. Orange dots show mean residual reaction time as a function of the binned normalized residual gain, with errorbars showing 95% confidence interval of this mean. Black regression line is best-fit linear regression of residual reaction time as a function of normalized residual gain, with shaded area showing 95% confidence interval of this regression. The regression slope is significantly negative for both monkeys ( *R* = −0. 14±0. 016, *p* < 10^−10^ for Monkey P; *R* = −0. 04±0. 015, *p* = 0. 007 for Monkey E).

While cross-validated model likelihood is generally robust for comparing predictive performance, it can still yield misleading conclusions when models differ in their computational complexity^52,53^. More flexible models may achieve better likelihood scores not because they better capture the underlying neural computation, but because they can more easily fit trial-by-trial variability, even if that variability reflects noise or incidental structure. This risk is especially relevant when comparing models with differing parameterizations, such as trial-specific variables or neuron-specific embeddings, which can lead to overfitting that is only partially penalized by cross-validation^34,54,55^. To avoid these risks, we used early stopping — evaluating models at the point of highest cross-validated likelihood, before overfitting began. We also tuned each model’s parameters separately for each dataset (see Methods). These steps helped ensure that differences in performance reflect meaningful differences between the models.

As a final measure of validity, we used synthetic data with known ground truth to assess whether overfitting could bias our results. We generated Poisson-sampled spike trains using the fitted models and applied our full model fitting and cross-validated likelihood comparison procedures to this synthetic data. This analysis confirmed that our model comparison reliably recovered the ground truth for each of the Slot, Switching, and Gain models (Fig. 4b). This result further supports the effectiveness of our modeling approach for testing hypotheses.

### Gain model predictions for behavioral errors and reaction times

Our model comparison showed that the Gain model of multi-object working memory is more consistent with neural data than the Slot and Switching models. A key feature of the Gain model is the recovery of per-trial gains, *w*_*k*_(*o*), associated with each object. These gains were distributed across the objects and did not sparsely select one object per trial (Extended Data Fig. 7). Consistent with extensive work in sensory^56–58^, association^22,23^, and frontal cortex^46,59,60^, we interpret these gains as trial-by-trial attentional modulations. Accordingly, we asked whether the neurally inferred gains could predict trial-by-trial accuracy and reaction time. Strikingly, performance was higher for objects with higher delay-phase gains (Fig. 4c), and residual gains were negatively correlated with residual reaction times (Fig. 4d). These behavioral relationships were weaker for the Slot and Switching models (Extended Data Fig. 8), suggesting that the Gain model’s latent variables better capture behaviorally relevant structure.

### Replication of the Results with the Ring Dataset

Having found evidence for the Gain model of multi-object working memory in our neural and behavioral data from the Triangle dataset, we next sought to replicate these results in the Ring dataset, an independent dataset with different stimulus configurations recorded in separate sessions (Fig. 1e). In the Ring dataset, object positions were sampled continuously around a circle to verify that the monkeys were not overfitting to the three discrete spatial positions in the Triangle dataset and that our modeling results were not an artifact of discrete positional sampling. Both monkeys performed well on the Ring dataset and exhibited reaction-time variability similar to the Triangle dataset (Fig. 5a–b; Extended Data Fig. 8a–b). We recorded 3,051 well-isolated neurons (1,712 from DMFC and 1,339 from FEF) across 22 neurophysiology sessions. These neurons showed a variety of spatial receptive fields in both DMFC and FEF (Fig. 5c; Extended Data Fig. 9c) and diverse selectivities to position and identity in delay-phase firing rates (Fig. 5d), similar to the Triangle dataset (Fig. 2b). We also found a positive relationship between signal correlation and noise correlation (Fig. 5e; Extended Data Fig. 9d), corroborating our findings in the Triangle dataset (Fig. 2d) and suggesting trial-by-trial attentional variability.

**Figure 5:**
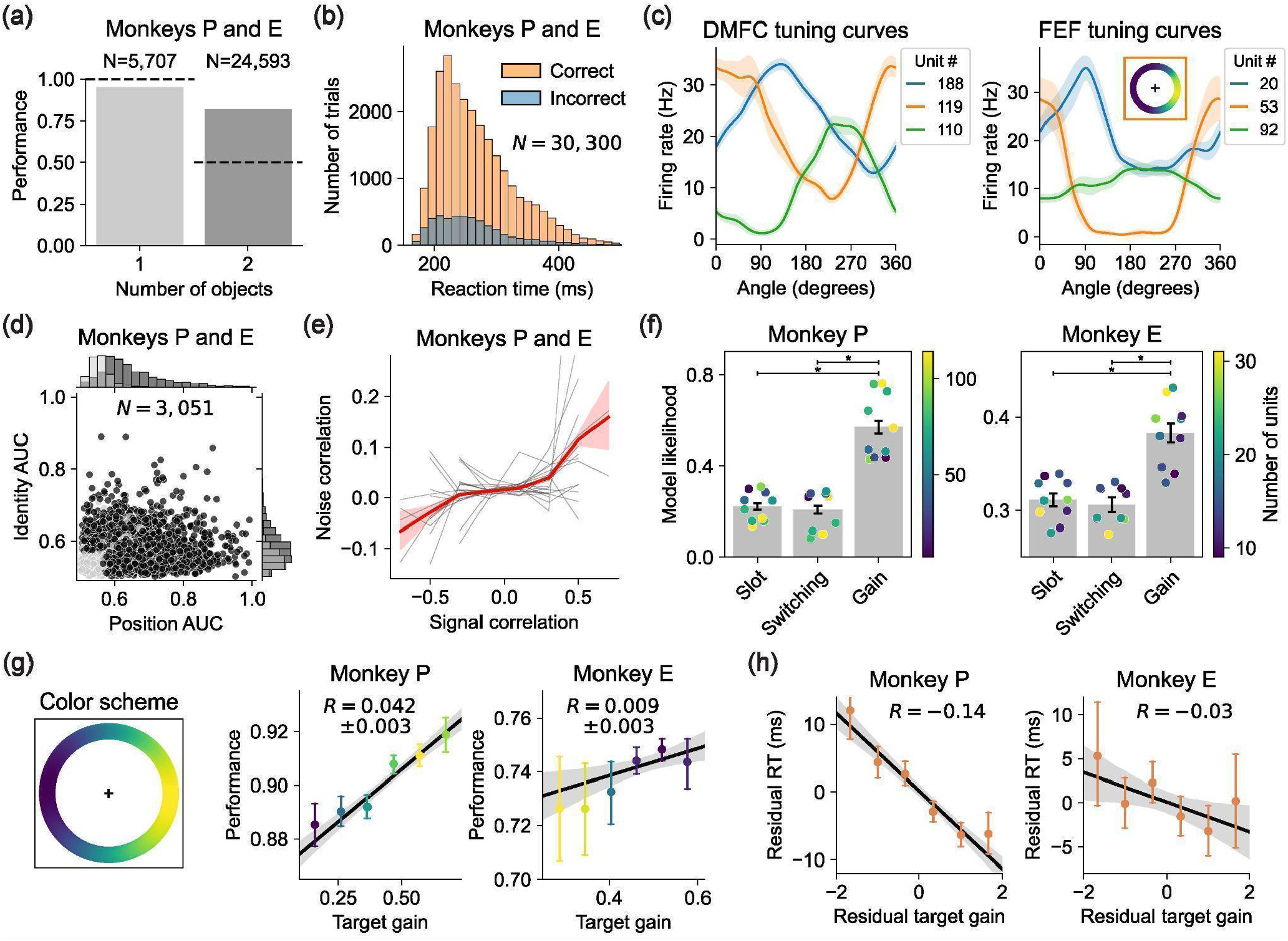
Ring dataset results. **(a)** Performance on the Ring dataset, aggregated over Monkeys P and E. Same plotting style as Fig. 1b. **(b)** Reaction time distribution on the Ring dataset, aggregated over Monkeys P and E. Same plotting style as Fig. 1c. **(c)** Angular tuning curves of three DMFC neurons (left) and three FEF neurons (right), from Monkey P session 2022-06-11. Each curve represents the delay-phase firing rate of one neuron as a function of stimulus object position, marginalizing over stimulus object identity. The errorbands show standard error of the mean. The inset in the right plot shows the angular tuning of unit 53 in the frame of reference of the display: For each position on the circle, the hue is proportional to the delay-phase firing rate for 1-object trials with an object at that position, smoothed by a Gaussian spatial kernel. **(d)** Position- and identity-selectivity of every recorded unit in the Ring dataset using area under the receiver operator characteristic (AUC) for 1-object trials. Same format at Fig. 2b, except both animals are included in the plot. **(e)** The relationship between noise and signal correlations for all pairs of simultaneously recorded neurons in the Ring dataset. Same format as in Fig. 2d. **(f)** Model likelihood for the Slot, Switching, and Gain models using the Ring dataset. Same format as in Fig. 4a. **(g)** Correlation between target gain and performance for 2-object trials from both monkeys in the Ring dataset, analogous to Fig. 4c. *Left:* Color scheme for hue as function of x-position of the target object. *Middle:* Performance (y-axis) as a function of gain of the target object (x-axis) for Monkey P. Black line shows regression over all trials, with shaded area showing 95% confidence interval of this regression. Regression is significantly positive (*R* = 0. 042±0. 003, *p* < 10^−10^). Colored dots show mean performance values for six evenly spaced target gain bins, with errorbars showing 95% confidence interval, using the color scheme on the left. *Right:* Same as middle plot except for Monkey E. Regression is significantly positive (*R* = 0. 01±0. 003, *p* = 0. 006). **(h)** Residual reaction time (y-axis) as a function of residual normalized target gain (x-axis) for correct 2-object trials on the Ring dataset for Monkey P (left) and Monkey E (right). Same format as Fig. 4d. The regression is significantly negative for both monkeys (*R* = −0. 13±0. 011, *p* < 10^−10^ for Monkey P, *R* = −0. 03±0. 012, *p* = 0. 018 for Monkey E).

All of our model formalizations apply to the Ring dataset without modification (see Methods). As in the Triangle dataset, the Gain model had higher model likelihood than the Slot and Switching models for both monkeys (Fig. 5f). Furthermore, the gain factors inferred by the Gain model again predicted both behavioral errors and reaction times (Fig. 5g–h).

## Discussion

Our work builds on a longstanding body of research on the computational basis of multi-object working memory in cognitive science and neuroscience^12,13,61^. Across these studies, three leading hypotheses have received sustained attention: the Slot hypothesis, the Switching hypothesis, and the Gain hypothesis. Each makes distinct claims about the neural mechanisms underlying working memory but has proven difficult to test directly, and as a result no consensus has emerged about which, if any, accurately reflects how the brain operates.

To address this question, we combined a large-scale neural dataset with a computational framework for formalizing and testing competing hypotheses. Three methodological components were especially important: formal encoding models, validation using synthetic data, and replication across independent datasets.

Our modeling framework formalizes hypotheses as computational constraints on neural activity and evaluates how well these constraints match neural data using encoding models. Each encoding model predicts the firing rate of each neuron on every trial and time point during the delay phase according to the assumptions of the corresponding hypothesis. This approach builds on a long tradition of using encoding models in neuroscience^62,63^ that affords quantitative precision, and direct, head-to-head comparisons (Extended Data Fig. 3c) avoiding ambiguous verbal descriptions^64^. Moreover, we validated our modeling framework using synthetic data, which is critical when evaluating complex statistical models^65–67^.

We initially studied one neural dataset, the Triangle dataset, performing all analyses including hypothesis testing and behavioral prediction. We then turned to a second, independently collected dataset — the Ring dataset — and found that our results replicated there. That our findings were not specific to a particular sample of neurons or stimulus configuration suggests that our conclusions generalize beyond a single experimental context. In both datasets, we found the Gain model to be most consistent with the neural data and providing the best explanation for behavioral accuracy and reaction times.

Our overall approach involved trade-offs and limitations that future research could address. First, to perform well-controlled high-density recording at the level of single neurons, we adopted the classical experimental paradigm of recording neural activity in awake, behaving monkeys in a laboratory setting. This experimental paradigm introduces certain limitations. For example, the use of virtual stimuli as opposed to real objects, long periods of training for the monkeys to learn the task, and the demand for gaze fixation may impact animals’ strategy and could thus affect the nature of neural computations.

Second, our modeling approach is limited by the hypotheses and assumptions therein. We focused on the Slot, Switching, and Gain hypotheses because of their prominence in the literature, but alternative accounts exist, including hierarchical models^68^, rate-distortion models^69^, gist-based models^70^, and hybrid models^71^. Furthermore, we evaluated all models using cross-validated Poisson likelihood. Although Poisson models are widely used, neural spiking statistics can deviate from Poisson assumptions^72,73^, and our comparisons may be sensitive to these assumptions. Future work may explore alternative assumptions, model metrics, and computational hypotheses. We have open-sourced our data and code to facilitate testing such alternatives.

Third, our results are specific to two cortical areas, DMFC and FEF. Other regions, including parietal and inferotemporal cortex^23,74^, are also involved in multi-object visual working memory and may use different representational formats. Applying our framework to additional areas will help clarify the anatomical specificity of our findings and the broader diversity of memory representations.

Lastly, our work has implications for several lines of research:

### Working memory capacity limitation

Unlike the Slot model, the Gain model does not by itself explain limited working-memory capacity. In principle, sufficiently high-dimensional noiseless embeddings would afford unlimited capacity. In practice, noise in neural activity likely creates ambiguity in stimulus representation, especially for objects with low gain. Trial-by-trial noise may perturb embeddings and cause memory failures. Inferring this noise alongside the latent variables of the Gain model may enable prediction of individual behavioral responses and clarify capacity limits. Future modeling work should test this prediction.

### Position-identity binding

A common error in our task occurs when the monkey reports the identity of an object at the wrong location (Extended Data Fig. 1), consistent with misbinding in working memory^32^. In our framework, the Gain model offers a natural account: objects with higher gain generate stronger population activity and may dominate retrieval^75,76^. When the cued identity is associated with a low-gain representation, the system may instead retrieve the location of a higher-gain object. This interpretation aligns with race-to-threshold models in which gain modulates evidence accumulation^77,78^ and is consistent with our finding that delay-phase gains predict reaction times. Future work should examine how gain-modulated traces interact with recall dynamics and shape binding performance^79^.

### Sequential multi-object working memory

Our results favor the Gain model for simultaneously presented stimuli, contrasting with recent findings supporting Slot-like subspaces during motor-sequence memorization^80,81^. A compelling interpretation of this discrepancy is that task demands shape the representational format of working memory. In those tasks, items were presented and recalled sequentially, whereas in ours, stimuli were presented simultaneously and only one item was cued for recall. This discrepancy suggests that working-memory formats are flexible and goal-sensitive rather than fixed buffers^82–84^. We therefore speculate that the same stimulus may yield different memory formats depending on what the brain plans to do with it. Exploring how task goals shape representational formats is an important direction for future work.

## Methods

### Open-Sourced Data and Code

We have open-sourced all of our data and code, as well as detailed instructions for reproducing our results. Visit https://github.com/jazlab/multi_object_memory_2025 for the instructions and code, and https://dandiarchive.org/dandiset/000620 for the data.

### Experimental procedures

All experimental procedures conformed to the guidelines of the National Institutes of Health and were approved by the Committee of Animal Care at the Massachusetts Institute of Technology. We performed all behavioral and neural experiments on two rhesus monkeys (Macaca mulatta). Monkey P was female, 10 years old, and weighed 8kg. Monkey E was male, 10 years old, and weighed 12kg. During all experiments the monkeys were seated comfortably in a primate chair and head-fixed in a dark room. The task was displayed on an Acer H236HL LCD monitor (23 inches diagonal size, 60 Hz refresh rate, 1920 by 1080 pixel resolution) positioned 22 inches away from the monkey’s eyes. The pupil position of the right eye was recorded at 1kHz by an infrared camera (Eyelink 1000, SR Research Ltd., Ontario, Canada). A mapping from pupil position in Eyelink camera coordinates to gaze point on the screen was calibrated at the beginning of every session. To do this calibration, the monkey was trained to fixate sequentially on each point in a 3×3 grid spanning the height and width of the display. The intervals of fixation were extracted and the eye positions during these intervals were used to fit a linear regression from pupil position to screen coordinate.

### Behavioral Task

The task was implemented in MOOG^85^ and MWorks (http://mworks-project.org). Below are details and parameters of each task phase:

#### Fixation phase

A square arena with sidelength 24° appeared on the screen. The arena was filled with grayscale pink noise with power spectrum density 1/f and normalized to have maximum luminance 0.25 times the full screen brightness. This background was randomly sampled each trial but unchanged within a trial and served to minimize afterimage effects that could be used for visual working memory. In the center of the arena was a white fixation cross with diameter 0.94°. If the gaze position was within 3.36° of the center of the fixation cross, the monkey was considered to be fixating. One of our monkeys had difficulty maintaining fixation, so we introduced a variable fixation cross size for both monkeys at all times the monkey was fixating: If the gaze position was further than 1.44° from the center of the fixation cross, the fixation cross grew in size linearly proportional to the gaze eccentricity passed through a 100ms boxcar filter. After holding fixation for 400ms, the task progressed to the stimulus phase.

#### Stimulus phase

Objects appeared with eccentricity 8.4° from the fixation point. Each object was a white circle of radius 2.06° with an image of a fruit occupying most of the circle. During physiology sessions, the fruit image was randomly sampled from a set of three images {apple, blueberry, orange}, though the animals were trained on a set of six images. The fixation cross remained through this phase. After 1000ms, the task progressed to the delay phase unless the monkey broke fixation.

#### Delay phase

The stimulus objects vanished, and the display looked exactly the same as during the fixation phase. After 1000ms, the task progressed to the cue phase unless the monkey broke fixation.

#### Cue phase

The cue was a white square with sidelength 4.12° presented at the center of the screen and occluding the fixation cross. Inside the square was rendered a random one of the fruits present in the stimulus of the trial at hand. After 180ms, the monkey was allowed to make a saccadic response, provided the fixation was maintained until that point.

#### Response phase

250ms after cue onset, a visual change occurred that helped indicate the monkey to respond. The cue became partially transparent (alpha 0.59) and a foveal window appeared. The foveal window was a circle with radius 2.4° that moved with the eye such that the center of this circle was the monkey’s gaze point. The foveal window effectively created a “hole” in the background noise, through which the monkey could see the objects hidden beneath. This was a useful tool during training, because it allowed the monkey to search for the correct stimulus object without using peripheral vision. The foveal window had a smooth boundary, with full background transparency within 1.2° and a linear radial gradient of opacity to full background opacity at 2.4°. The foveal window did not allow the monkey to view any stimulus objects after the cue onset before making a response saccade, hence did not allow the monkey to cheat on the task. The foveal window also did not give rise to any unusual eye movement behaviors. The monkeys made saccadic responses with normal eye velocity and eccentricity. The endpoint of a response saccade was detected if the gaze position was no longer within 3.36° of the center of the screen and the gaze velocity was below 7.2 degrees per second. In practice, these parameters detected saccadic responses very reliably for both monkeys.

#### Feedback phase

After the monkey responded, all objects were revealed with partial opacity (alpha 0.59) for visual feedback. This feedback lasted 500ms for correct trials and 1000ms for incorrect trials. During the feedback phase the monkey received a water or juice reward for correct trials. A response was deemed correct if it lay within 2.4° of the center of the target object. The reward was delivered via a tube next to the monkey’s mouth, and the flow of reward was controlled by a solenoid. The reward scaled with trial difficulty: The monkey received 100% of the reward duration for correct 3-object trials, 70% for correct 2-object trials, and 40% for correct 1-object trials. The reward also scaled with accuracy, with maximum reward for a response position exactly the same as the center of the target object. Reward scaled linearly down to zero if the response was more than 2.4° from the center of the target object. Upon a correct response, in addition to reward, the monkey received an auditory cue (high-pitched ding). Upon an incorrect response, the monkey received a different auditory cue (low-pitched honk).

#### Inter-trial interval

A gray screen (luminance 0.2 times the maximum brightness of the screen) was presented for 500ms after correct trials and 1000ms after incorrect trials before the fixation phase began for the next trial.

If the monkey’s gaze position was greater than 3.36° of the center of the screen at any point until the response phase, the monkey was considered to have broken fixation. If the monkey broke fixation during the fixation phase of the task, the trial did not restart and the task waited for the monkey to regain fixation. If the monkey broke fixation during the stimulus, delay, or cue phase, the trial aborted. In this case, as soon as fixation was broken, the arena turned into a maroon square (RGB value [102, 51, 51]) for 1000ms, followed by a 1000ms ITI, followed by the onset of the next trial. Simultaneously, the monkey got auditory feedback (same low-pitched honk as for incorrect trials).

For the Triangle dataset, to encourage the monkeys to have effortful behavior for difficult 3-object trials we used a performance-conditioned block structure: The task presented 3-object trials until the monkey got 10 of those correct, then 2-object trials until the monkey got 10 of those correct, then 1-object trials until the monkey got 3 of those correct, then back to 3-object trials, and so on. For the 3-object trials, we sampled each trial as follows: With probability 0.9, the trial was sampled from one of the 6 possible 3-object triangle stimuli (the different combinations of object identities in the three vertices of the triangle); with probability 0.1, the three objects’ positions were sampled randomly with eccentricity 8.4° subject to the constraint that no two objects were within 9.2° of each other. We added these trials with random object positions to prevent the monkey from overfitting to the triangle positions. For the 2-object trials, we sampled each trial as follows: With probability 0.9, the trial was sampled from one of the 18 possible 2-object triangle stimuli (the different combinations of object identities in each pair of vertices of the triangle); with probability 0.1, the two objects’ positions were randomly sampled with eccentricity 8.4° subject to the constraint that no two objects were within 9.2° of each other. For the 1-object trials, we sampled each trial from one of the 9 possible 1-object trial stimuli (different combinations of object identity and location). We restricted all analysis and modeling to the trials with positions on the triangle. The triangle positions formed the vertices of an equilateral triangle with centroid at the center of the screen. The triangle was oriented such that one vertex was directly to the right of center and the other two vertices were on the left side of the screen (one in the top-left quadrant and the other in the bottom-left quadrant).

For the Ring dataset, we randomly interleaved 1-object trials with 2-object trials. For each trial, with probability 0.75 we delivered a 2-object stimulus in which the two objects’ positions were randomly sampled with eccentricity 8.4° subject to the constraint that no two objects were within 9.2° of each other. With probability 0.2 we delivered a random 1-object stimulus in which one object was randomly uniformly sampled with eccentricity 8.4°. With probability 0.05 we delivered a left/right 2-object stimulus in which one object was 8.4° directly to the left of fixation and the other was 8.4° directly to the right of fixation. We did this to make possible analyses that rely on trial averaging over replicated conditions, though did not end up doing such analyses. In all conditions the object identities were uniformly sampled and the cued object was randomized. We used all conditions for analysis.

### Neurophysiology

Each monkey underwent a first surgery under general anesthesia to attach three titanium headpins to the skull for head fixation and a second surgery to implant a medial chamber of polyether ether ketone for neurophysiology recording. A craniotomy was made over DMFC in each monkey. We recorded simultaneously from right-hemisphere dorsomedial frontal cortex (DMFC) and left-hemisphere frontal eye field in both monkeys. DMFC includes supplementary eye field (SEF), presupplementary motor area (Pre-SMA), and dorsal supplementary motor area (SMAd)^66,86,87^. We recorded from a 4mm x 6mm patch of superficial DMFC in each monkey, randomly sampling recording sites within this patch during each session. In frontal eye field we recorded from the lateral bank of the superior arcuate sulcus and the anterior bank of the genu of the arcuate sulcus, randomly sampling recording sites from a 5mm x 8mm patch of cortex during each session. See Extended Data Fig. 10a for a visualization of recording sites.

We recorded from DMFC using non-reinforced 10mm Neuropixels 1.0 probes^88^. Because these probes are very short and fragile, we developed a custom guide tube system (Extended Data Fig. 10b). This guide tube system allowed us to penetrate the primate dura with a very short (2mm) tip of the guide tube and insert the Neuropixels probe through the guide tube into DMFC via a micromanipulator, while controlling the stereotaxic coordinates of the insertion.

We recorded from FEF using Plexon V-probes with 64 channels and 50 micron inter-contact spacing. We used V-probes to record from FEF because both monkeys had pre-existing chambers centered over the frontal midline for DMFC and did not lie over FEF, so we had to approach FEF through an angled trajectory that was too long for 10mm Neuropixels probes.

We used a custom angled grid and guide tube system for FEF recordings that was compatible with simultaneous DMFC recording.

We recorded Neuropixels data at 30kHz frequency with SpikeGLX software and V-probe data at 30kHz frequency with OpenEphys software.

For the Triangle dataset, we recorded 24 total sessions (10 from Monkey P and 14 from Monkey E). For the Ring dataset, we recorded 22 total sessions (10 from Monkey P and 12 from Monkey E).

## Data Processing

Neural data were temporally synchronized to behavioral data by a linear regression based on a sync pulse sent at the beginning of every trial, and the timing of task events on the monitor was adjusted by a display delay measured via a photodiode in the corner of the monitor.

Neural data was then spike sorted using Kilosort 2.5^89^. For V-probe data, spike sorting results were curated manually with a spike-sorting curation gui (Phy), through which we did splitting, merging, and labeling of the units. For Neuropixels data, we performed motion estimation and correction using MEDiCINe and SpikeInterface^90,91^. The quantity of units from the Neuropixels data was prohibitively large for manual merging and splitting, so we used a custom automated pipeline to do splitting based on unit statistics, then manually labeled each unit to identify putative single-units and multi-units. Our pipeline also detected and excluded duplicated units, which have the potential to give spurious modeling results. Broadly, we were extremely conservative with our spike sorting: We thoroughly examined a variety of statistics and plots for every unit, only labeled units as single neurons if they were very clearly single neurons, and liberally classified as noise anything that did not have a clear neuron-like waveform.

After spike sorting, we identified for each recorded unit an interval of time for which the unit was stable by running a filter on firing rate, thresholding the derivative of firing rate over time to identify points where the neuron was lost or gained, and validating manually. For many neurons the identified interval of stable recording was the entire session, but some neurons were lost or gained partway through the session. We excluded all spikes outside of this stable range. We also excluded from analysis all neurons that were not stable for at least 500 trials, and all neurons with mean firing rate less than 1Hz during the delay phase of the task.

### Receiver-Operator Characteristics

To compute the position AUC value of a unit *n*, we first computed the mean firing rate *r*(*k*) of unit *n* during the delay phase for each 1-object trial *k*. We then assigned each 1-object trial *k* a label in {1, 2, 3} based on where the object was. For the Triangle dataset this was which of the three locations the object occupied. For the Ring dataset this was which of the three locations in the Triangle dataset was nearest to the object (i.e. dividing space into three equal angular sections). We then computed the receiver-operator function for locations 1 and 2. To do this, let *g*_*i*_ (*x*) = #[*r*(*k*_*i*_) < *x*]/*N*_*i*_ where *N*_*i*_ is the number of trials with label *i*, so *g*_*i*_ (*x*) is the fraction of trials with label *i* that have mean delay-phase firing rate less than *x*. Now note that plotting *g*_1_ (*x*) on the x-axis and *g*_2_ (*x*) on the y-axis will trace a path that monotonically increases in both x and y from the (0, 0) to (1, 1). Let *AUC*_1,2_ be the area under this path, namely between this path and the x-axis interval [0, 1]. Thus by construction *AUC*_1,2_ must be between 0 and 1. Note that *AUC*_1,2_ measures how much the firing rate prefers location-2 to location-1 trials. For example, if *r*(*k*_2_) > *r*(*k*_1_) for all trials then *AUC*_1,2_ = 1, whereas if *r*(*k*_2_) < *r*(*k*_1_) for all trials then *AUC*_1,2_ = 0. Lastly, we define *AUC*_(1,2)_ = *max*[*AUC*_1,2_, *AUC*_2,1_], so *AUC*_(1,2)_ > 0. 5 and measures how discriminable location-1 and location-2 trials are based on the delay-phase firing rate of unit *n*. To compute the p-value of *AUC*_(1,2)_, we randomly shuffled the trial labels 1,000 times and computed the resulting *AUC*_(1,2)_ for each shuffle. We then let the p-value *p*_(1,2)_ be the fraction of these shuffled null *AUC*_(1,2)_ values that are greater than the empirical *AUC*_(1,2)_. Analogously, we computed *AUC*_(1,3)_ and *AUC*_(2,3)_, the AUC values for the other pairs of locations, and their associated p-values. If none of *AUC*_(1,2)_, *AUC*_(1,3)_, and *AUC*_(2,3)_ had p-value less than 0.05, then we let the position AUC value be *AUC*_*position*_ = *max*(*AUC*_(1,2)_, *AUC*_(2,3)_, *AUC*_(1,3)_). Otherwise, we let *AUC*_*position*_ = *max*([*AUC*_(*i,j*)_| *p*_(*i,j*)_ < 0. 05]). To compute the identity AUC value *AUC*_*identity*_, we performed exactly the same procedure except labeled the 1-object trials by object identity instead of position. We similarly had three labels, since our stimuli during physiology had three possible object identities (apple, blueberry, and orange). The x-axis and y-axis of Fig. 2b are *AUC*_*position*_ and *AUC*_*identity*_, respectively.

### Noise Correlations

The plots in Fig. 2c are marginalized over object identities. However, the correlations in Fig. 2c are similar when conditionalizing on object identity (Extended Data Fig. 4), so the correlations are not due to identity tuning.

For Fig. 2d, we computed the signal correlation and noise correlation for each pair of simultaneously recorded units. To compute the signal correlation for units *n*_1_ and *n*_2_, we considered every possible 1-object condition in the Triangle dataset. There were 9 such conditions (three identities times three locations). If either unit *n*_1_ or unit *n*_2_ was not recorded for at least 3 trials on each of these 9 conditions, we discarded this pair of units for lack of data. We then computed the correlation coefficient between the mean delay-phase firing rates of unit *n*_1_ and the mean delay-phase firing rates of unit *n*_2_ across these 9 conditions. This correlation coefficient is the signal correlation between units *n*_1_ and *n*_2_ and is the x-axis of the plots in Fig. 2d.

To compute the noise correlation of units *n*_1_ and *n*_2_, we considered every possible 2-object condition in the Triangle dataset. There were 18 such conditions (three identity pairs times three location pairs times two permutations). For each such condition, we computed the correlation of delay-phase firing rates across trials for which the neurons were simultaneously recorded, excluding conditions for which we did not have at least 3 trials with simultaneous recording of the units. The mean of these correlation coefficients is the noise correlation between units *n*_1_ and *n*_2_, which is the y-axis of the plots in Fig. 2d.

### Modeling

We parameterized all variables of all models in a differentiable manner, allowing fitting by gradient descent. For each model, we parameterized *b*(*n, t*) by a multilayer perceptron (MLP) with output size *N*_*units*_, hidden sizes [512, 1024, 512], and input size 5. Given an input time *t* in units of seconds, we computed 4 sinusoid features to make the MLP input a 5-vector [*t, sin*(*t*), *sin*(2*t*), *sin*(4*t*), *sin*(8*t*)], and let *b*(*n, t*) be the *n*’th component of the MLP’s output. The sinusoid input features helped the model more easily learn temporal modulations.

We parameterized each embedding function (*f*_*o*_ (*n, t*) for the Gain and Switching models and {*f*_*s,o*_ (*n, t*)}_*s*∈{1,2}_ for the Slot model) by an MLP with output size *N*_*units*_, hidden sizes [512, 1024, 512], and input size 18. Given an input time *t* in units of seconds and an object with position (*x, y*) in units of display sidelength and identity α, we computed feature vectors

*t*_*features*_ = [*t, sin*(*t*), *sin*(2*t*), *sin*(4*t*), *sin*(8*t*)],

*x*_*features*_ = [*x, sin*(*x*), *sin*(2*x*), *sin*(4*x*), *sin*(8*x*)],

*y*_*features*_ = [*y, sin*(*y*), *sin*(2*y*), *sin*(4*y*), *sin*(8*y*)],

α_*features*_ = [α = “*apple*”, α = “*blueberry*”, α = “*orange*”] (in other words α_*features*_ was a one-hot code for the object identity). We then let the input to the embedding MLP be the concatenation of *t*_*features*_, *x*_*features*_, *y*_*features*_, and α_*features*_. The sinusoid features of position and time helped the model more efficiently learn spatial and temporal modulations.

For all models, both the baseline function *b* and the object embedding function(s) *f* were at risk for overfitting. To avoid this, we implicitly pressured them to vary smoothly in time (and in space for the object embedding functions) by adding noise during training. Specifically, during training, to every input time *t* we added random Gaussian noise, using instead *t* + *N*(0, κ) as the input to the temporal feature vector. We tuned the temporal noise scale κ on a per-session and per-model basis to minimize overfitting, as described below. For the object embedding functions, during training we added independent Gaussian noise with standard deviation 0.05 to both *x* and *y* components of the object’s position. The positional noise was largely irrelevant for the Triangle dataset, but did reduce overfitting for the Ring dataset where object positions varied continuously.

### Slot Model

The Slot model has two discrete functions: The object-slot assignment function σ_*k*_ (*s*) and the allocation of which neurons belong to which slots. Let *I*(*n*) = *s*∈[1, …, *N*_*slots*_] denote an indicator function allocating each neuron *n* to a slot. *I*(*n*) and σ_*k*_ (*s*) are discrete functions. Hence it is not immediately clear how to optimize *I*(*n*) and σ_*k*_ (*s*) using gradient descent. To solve this problem, we used a loss-weighting technique to fit continuous approximations of these parameters in a way that in principle recovers their discrete optima. To do this, we parameterized *I*(*n*) by a continuous matrix of size [*N*_*units*_, *N*_*slots*_], and we parameterized σ by a continuous matrix *A* of size [*N*_*trials*_, *N*_*assignments*_], where *N*_*assignments*_ is the maximum number of possible functions σ_*k*_. Through training, *I* and *A* are implicitly pressured to become one-hot vectors for each unit and trial, respectively. Through hyperparameter sweeping, we found that *N*_*slots*_= 2 provided the best cross-validated performance on both the Triangle and Ring dataset for both monkeys. Given this, *N*_*assignments*_ is 6 for the Triangle dataset (since it has a maximum of 3 objects) and 2 for the Ring dataset (since it has a maximum of 2 objects). To formulate the objective function for the Slot model, we computed the log likelihood loss on every trial for each unit, slot, and assignment:

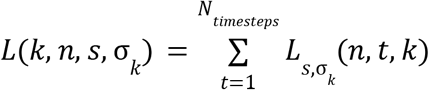

where

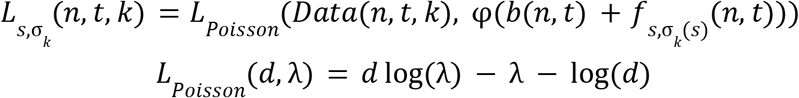

Thus *L*(*k, n, s*, σ_*k*_) is the loss on trial *k* for assignment σ_*k*_ if unit *n* belongs to slot *s*. We then compute the total loss

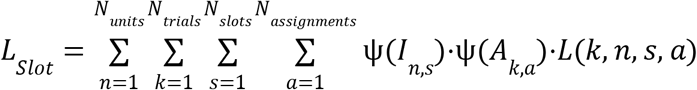

where

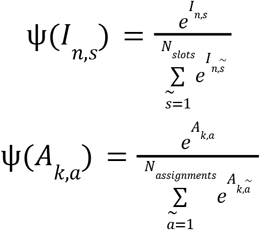

are softmax functions of *I*_*n*,−_ and *A*_*k*,−_ and we have indexed each assignment σ_*k*_ using an index *a*∈[1, …, *N*_*assignments*_] for notational convenience. Intuitively, the loss function *L*_*Slot*_ sums the losses over each allocation of unit to slot and each assignment of slot to object, weighted by the logits *I*_*n*,−_ and *A*_*k*,−_. This objective function implicitly pressures *I* and *A* to be one-hot, because they are pressured to put all their weight on the slot and assignment that gives the lowest loss. For evaluation, we discretize *I* and *A* by applying an argmax function along the last dimension.

To tune the parameter *N*_*slots*_, for each monkey and dataset we selected 4 random sessions and explored values *N*_*slots*_ ∈[1, 2, 3, 4] for 2 random train/test splits for each of these sessions. In all cases we found that on average, cross-validated log likelihood was best for *N*_*slots*_ = 2. We then used *N*_*slots*_ = 2 for all remaining results.

### General Slot Model

As mentioned in the main text, allowing the slots to consist of arbitrary subspaces (not restricted to disjoint sets of neurons) did not qualitatively impact our results (Extended Data Fig. 5). To formalize this more general variant of the Slot model, we modified the formalization above by removing the neuron-slot indicator function *I*(*n*). This simplifies the loss function. We computed the log likelihood loss on every trial for each slot and assignment:

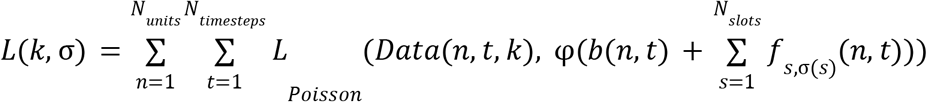

Thus *L*(*k*, σ_*k*_) is the conditional loss for assignment σ_*k*_. We then computed the total loss

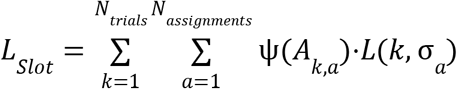

where σ_*a*_ is an enumeration of the *N*_*assignments*_ assignments and

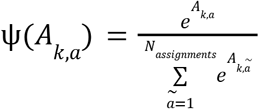

is a softmax function of *A*_*k*_. For evaluation, we binarized Ψ(*A*_*k,a*_) to be one-hot for each *k*.

We used the same approach as above to tune the parameter *N*_*slots*_ = 2 was best for both monkeys. *N*_*slots*_, and also found that

### Switching Model

In the Switching model, ρ_*k*_ (*t*) is a discrete variable. We fit ρ_*k*_ (*t*) using a similar approach to fitting the discrete variables in the Slot model. We parameterized ρ_*k*_ by a matrix *R* of shape [*N*_*trials*_, *N*_*timesteps*_, 1 + *N*_*objects*_] where *N*_*objects*_ is the maximum number of objects in a trial. This matrix *R* represents logits on the likelihood of ρ_*k*_ (*t*) being assigned to each of the objects (or the empty token). We then smoothed *R* through time with a triangular kernel with 200ms support. This smoothing forces the model to take at least 100ms to completely switch from having full weight for one object to having full weight for another object. In practice we found our results qualitatively insensitive to this parameter. Let *Q* denote this smoothed version of *R*. We computed the loss function for each trial and timestep per object:

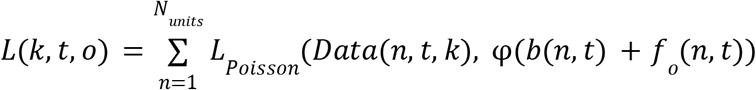

where *o*∈*Objects*(*k*) + [*empty*]. We then defined the total loss as

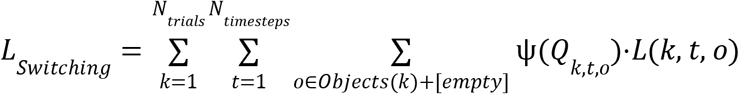

where

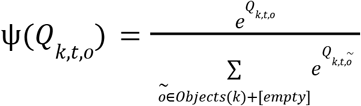

is a softmax function on *Q*_*k,t*_. Intuitively, the loss function *L*_*Switching*_ sums the losses over each possible value of the switching latent, weighted by the logits *Q*. This objective function implicitly pressures *Q*_*k,t*_ to be a one-hot vector putting all of its weight on the object that gives the lowest loss. For evaluation, we discretize *Q* by applying an argmax function along the last dimension to get a discrete switching variable ρ. To make *Q* well-defined for trials with fewer than *N*_*objects*_ objects, for each trial we mask *Q* inside the softmax function Ψ to enforce the resulting logit takes value zero for excess dimensions.

### Gain Model

We parameterized *w*_*k*_ (*o*) in the Gain model by a matrix *W* of shape [*N*_*trials*_, *N*_*objects*_]. To bound the gain factor, we use a sigmoid function 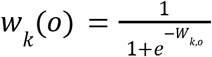. We chose to bound the gain factor to [0,1] because this is similar to normalization models of attention in prior work ^7,11^, but we found qualitatively similar results without the upper bound of 1. Since *w*_*k*_ (*o*) is continuous, we computed the loss *L* directly as a function of *w*_*k*_ (*o*) and backpropagate gradients through all variables.

### Model-Fitting

Running our modeling pipeline on all of our data would have been prohibitively computationally expensive, so we included only units that had significant task modulation. Specifically, we only included units that, for either position AUC or identity AUC in the delay phase, had both AUC value greater than 0.65 and AUC p-value less than 0.05.

We implemented all models in PyTorch. For all models, we optimized all parameters with Adam optimizer with learning rate 0.003 and a gradient clipping value of 1. For the first 1000 gradient steps, we only optimized the per-unit parameters (*b* and *f* in the model formulations), not the per-trial parameters. We did this because we found it to reduce overfitting: Giving the per-unit parameters *b* and *f* a head-start effectively served to initialize them in a reasonable regime and helped avoid the per-trial parameters learning based off of nonsensible unit embeddings. Another way to see is is because the per-trial parameters are very near to the loss function in the models, they are likely to fit too fast and disrupt the optimization of *b* and *f* unless *b* and *f* are already in a reasonable regime. We verified optimization was working well by (i) ensuring that the fitted parameters recovered the ground truth for our synthetic datasets, and (ii) ensuring that for our neural data the fitted parameters were consistent across random initializations and train/test splits.

For all main results, we fit and evaluated each model on 10 different train/test splits for each session. For each train/test split, we assigned each trial of each unit to the training set with probability 0.8 and to the test set with probability 0.2. We performed this train/test split on a per-unit-trial basis because all of our models have per-unit parameters and per-trial parameters, so the training data must include activity from some units for each trial and activity from some trials for each unit. The number of gradient steps we trained was a hyperparameter that we tuned on a per-session basis to maximize cross-validated likelihood (see below), like the temporal smoothness κ. We did this to help reduce overfitting. For all test-set evaluations we set κ to zero.

### Tuning the Temporal Smoothness and Training Steps

For each model, for every session we independently tuned both the embedding temporal smoothness κ and the number of training gradient steps to maximize cross-validated log likelihood. We did this by sweeping κ from 0 to 200ms in increments of 20ms for two random train/test splits and training each for 10000 gradient steps. We then considered the sum of test-set log likelihoods over these two runs and took the value of κ and the training step that maximized this. We used the resulting κ and the resulting training step for the main results.

We tuned κ and the training step independently per model because each model has a different computational complexity and hence may have a different optimal κ and training step for cross-validated log likelihood. We tuned κ and the training step independently per session because different sessions have different signal-to-noise and hence different optimal values of κ for cross-validated log likelihood. In particular, sessions with more simultaneously recorded units tend to have a lower optimal value of κ.

We believe this tuning procedure treated all models fairly and gave each model the best chance at explaining the data. We found that for most sessions, cross-validated likelihood was maximized for a value of κ somewhere in the middle of our sweep. Furthermore, this optimal value of κ was broadly consistent across models. See Extended Data Fig. 11 for visualizations of the test loss landscape as a function of κ and training steps for a typical session.

### Model Comparison

In Fig. 4a, we computed the test-set model likelihood for each model on each session of neural data. Recall that for each session of data, we fit all models independently on 10 train/test splits. For each of these splits, we then computed an estimate of the model likelihood *P*(*model* | *data*). To compute this estimate, we performed a bootstrapping procedure: For 1000 bootstraps, we drew a sample of 100 test neuron-trial pairs and computed the model likelihood on those, then averaged this likelihood over the bootstraps. We did this instead of computing the total likelihood over all test datapoints for two reasons. First, it allowed the likelihood for a session to not be dependent a priori on the number of neurons or trials for that session. Second, because we averaged over samples of only 100 neuron-trial pairs, this process gave likelihood values that were not as extreme as taking the likelihood over the entire test set, hence allowing the model comparison to be more visually apparent.

### Validation Using Synthetic Data

To generate the results in Fig. 4b, we considered 10 sessions of data: The five sessions with the most units from each monkey. We used a subset of sessions because tuning the temporal smoothness parameter κ was computationally very expensive. For each of these sessions we selected one train/test split and considered each trained model for that common train/test split. For each of the models, we took the state of the model at the training step with maximum cross-validated log likelihood (the one contributing to the results in Fig. 4a). From that model instance we sampled spiking data for every timestep, trial, and neuron under a Poisson assumption (and setting κ to zero). We saved these datasets as synthetic datasets. Intuitively, these are synthetic datasets that best match the neural data under the constraint imposed by each hypothesis.

We then treated these datasets as neural data and applied our whole modeling pipeline to them: We tuned κ and the stopping step independently for each session for each model, then fit all models on each of the synthetic datasets for two new train/test splits. The goal of this exercise was to determine whether, if the neural data had in fact come from one of our hypotheses, we would have recovered that hypothesis through our modeling procedure. Our model comparison metric was exactly the same bootstrapped model likelihood that we used for comparing models on our neural data, except we used 300 neuron-trial pairs for bootstrapping. This does not qualitatively affect the results, just makes the differences in model likelihoods more apparent, which is useful given the relatively lower variance of the synthetic data from the model data due to temporal smoothing and lack of spike timing effects.

### Ring Dataset Tuning Curves

In Fig. 5a, we computed the smoothed bootstrapped delay-phase firing rate for each neuron as a function of stimulus object angle. To do this, we computed 256 evenly spaced points on the ring. For each point, we computed the bootstrapped smoothed firing rate. To do this, consider a point *p* with position (*x, y*). We performed 100 bootstraps, where for each bootstrap we sampled 100 1-object trials with replacement. We then let the firing rate be 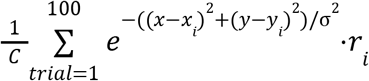, where (*x*_*i*_, *y*_*i*_) is the object position on trial *i, r*_*i*_ is the firing rate on trial *i*, σ is 0.1 times the sidelength of the display arena (namely, 2.4° of visual angle), and 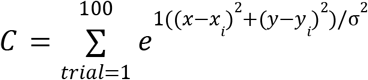 is a coefficient normalization factor. Intuitively, this is smoothing with a spatial 2-dimensional Gaussian kernel.

### Ring Dataset Signal and Noise Correlations

In Fig. 5c, we computed signal and noise correlations for pairs of units in the Ring dataset. Because the Ring dataset had continuously sampled stimuli, we could not use the same procedure for computing signal and noise correlations as for the Triangle dataset. Instead, we first needed to establish a metric of stimulus similarity across trials. To do this, we embedded each trial stimulus into a 6-dimensional space, where the axes were [apple-x, apple-y, blueberry-x, blueberry-y, orange-x, orange-y]. Here “fruit-x” is the x-position of the “fruit” object in the stimulus (0 if that fruit is not present), and “fruit-y” is the y-position of the “fruit” object in the stimulus (0 if that fruit is not present). In this 6-dimensional trial embedding space we used Euclidean distance between points as a measure of stimulus similarity across trials.

Given this, to compute the signal correlation for a pair of units, we considered 1-object trials for which both units were stably recorded. For each of these trials, we took the delay-phase firing rate of the first unit as one coordinate and the delay-phase firing rate of the second unit on the nearest-neighbor trial in the trial embedding space as a second coordinate. We then took the correlation of resulting datapoints as the signal correlation of the units. We did this nearest-neighbor procedure to ensure that our computation of signal correlation used firing rates from the two units on different trials, hence was not influenced by noise correlations.

To compute the noise correlation for a pair of units, we considered 2-object trials for which both units were stably recorded. For each such trial, we computed a noise correlation sample by finding the 10 nearest-neighbor trials in the trial embedding space and computing the (de-meaned) correlation of the firing rates of the units on these trials. Intuitively, these 10 trials all have extremely similar stimuli, so computing the correlation of the units on these trials is as close as we can get to a noise correlation for this dataset.

### Ring Dataset Normalized Target Gain

In Figure 5e and 5f, the normalized target gain (x-axis) is conditionalized on identity and position and normalized on a per-session basis to have variance 1. To compute this normalized target gain for a given trial, we take the target object gain factor, subtract the mean gain factor over all trials of objects with the same identity as the target object and nearby position to the target object (within 90° angle), then scale the resulting distribution of the result on a per-session basis to have variance 1. By conditionalizing on both target identity and target position in this way, we conclude that the resulting correlations must arise from trial-by-trial variance and are not explained by biases in attention over object position or identity.

### Decoding Analyses

The results in Extended Data Figure 3b show that decoding analyses are insufficient to determine whether a multi-object neural representation engages orthogonal slots in neural activity space. In contrast, with reasonably large numbers of simultaneously recorded neurons (in this case ~100), the curse of dimensionality causes almost every pair of axes in neural activity space to look orthogonal, regardless of the underlying representation structure they come from.

To show this, we focused on 2-object conditions in both the Triangle and Ring datasets, and built a 2-object linear decoder model that decoded the features of both objects from neural firing rates on single trials. To formalize this, first consider one Triangle dataset session. Let *N*_*units*_ be the number of simultaneously recorded units and *N*_*trials*_ be the number of 2-object trials for which all of these units were stably recorded. The 2-object decoding model then consists of the following 5 variables:

*W*_0_: Matrix of shape [*N*_*units*_, 3]

*B*_0_: Vector of length 3

*W*_1_: Matrix of shape [*N*_*units*_, 3]

*B*_1_: Vector of length 3

σ: Vector of length *N*_*trials*_

The variables *W*_0_, *B*_0_ parameterize one linear decoder from the space of neurons to the space of one-hot vectors coding object position (for these analyses we marginalize over identity, since identity was poorly decodable on single trials). The variables *W*_1_, *B*_1_ parameterize a second linear decoder. The variable σ is a logit constrained to [0, 1] by a sigmoid function, and σ determines which decoder to apply to which object on each trial. We implicitly pressure σ to be binary using the same loss-weighting technique as used for the Slot and Switching models. Specifically, we fit all of these variables simultaneously with gradient descent using the following loss function:

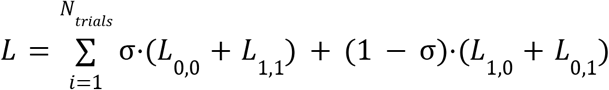

Where *L*_*i,j*_ is the error of the result of decoder *i* with respect to the one-hot position of object *j*. In practice we used mean squared error, though the results were qualitatively the same for binary cross entropy error.

After fitting this model, we then evaluated whether the weights *W*_0_ and *W*_1_ were orthogonal. Specifically, for each of the three locations, we measured the cosine angle between the corresponding columns in *W*_0_ and *W*_1_. That is shown on the y-axis of Extended Data Figure 3b. Intuitively, one might expect a resulting angle of 90 degrees to imply that the neural state space has orthogonal subspaces, each with the capacity to represent one object, and each encoding a different object on every trial, like the Slot hypothesis. However, we found that this orthogonality resulted even on synthetic data that was specifically not Slot-like. In other words, orthogonality of these decoding subspaces is not diagnostic of object slots. This is due to the curse of dimensionality: In high-dimensional space (e.g. neural state space), a randomly sampled pair of vectors is highly likely to be orthogonal. So even in a representation that doesn’t have slots, the 2-object decoder model will find some symmetry-breaking feature to determine how to assign objects to decoders (in practice, we found this feature to be a relationship between the object positions), and because of that symmetry breaking, the decoders will effectively have different training distributions, causing their subspaces to diverge in neural state space, hence be orthogonal by the curse of dimensionality.

For the Receptive Field synthetic dataset, we simulated Gaussian receptive fields randomly distributed in the display arena. We then let the response of each neuron be the sum of the contributions of the two objects, given the neuron’s receptive field. We normalized this response to have the same mean and variance across trials as the real data, on a per-neuron basis. For the Orthogonal dataset, we took two independent copies of the Receptive Field model (each with half the total number of neurons) and aggregated their resulting neurons (then re-normalized to have the same mean and variance as the real data). Thus in the Orthogonal dataset, each object is encoded by a disjoint set of units. The assignment of which object went to which subspace was randomized per trial. We verified that the 2-object decoding model recovered the ground truth subspaces accurately.

## Author contributions

N.W., J.T., and M.J. designed the task. N.W. and J.G. collected the data. N.W. did the data processing and implemented the analyses. N.W. and M.J. developed the models. N.W., J.G., J.T., and M.J. wrote the manuscript.

## Open-Sourced Data and Code

We have open-sourced all of our data and code, as well as instructions for reproducing our results and demo scripts for becoming familiar with our data. For the code, instructions, and demo scripts, visit https://github.com/jazlab/multi_object_memory_2025. For our data, visit https://dandiarchive.org/dandiset/000620.

## Acknowledgments

N.W. is supported by the NSF. J.T. is supported by the Siegel Family Quest for Intelligence at MIT, ANR Science of AI program, and SCGB. M.J. is supported by the Simons Foundation, HHMI, and the McGovern Institute. We thank Rishi Rajalingham, Sujay Neupane, Alexandra Ferguson, and Hansem Sohn for their guidance in setting up the experimental rigs and developing our neurophysiology methods. We thank the CatalystNeuro team including Cody Baker, Luiz Tauffer, Felix Pei, Alessio Buccino, and Ben Dichter for their help open-sourcing our data. We thank Michael Visconti for fabricating much of our experimental hardware.

## Extended Data Figures

**Extended Data Figure 1:**
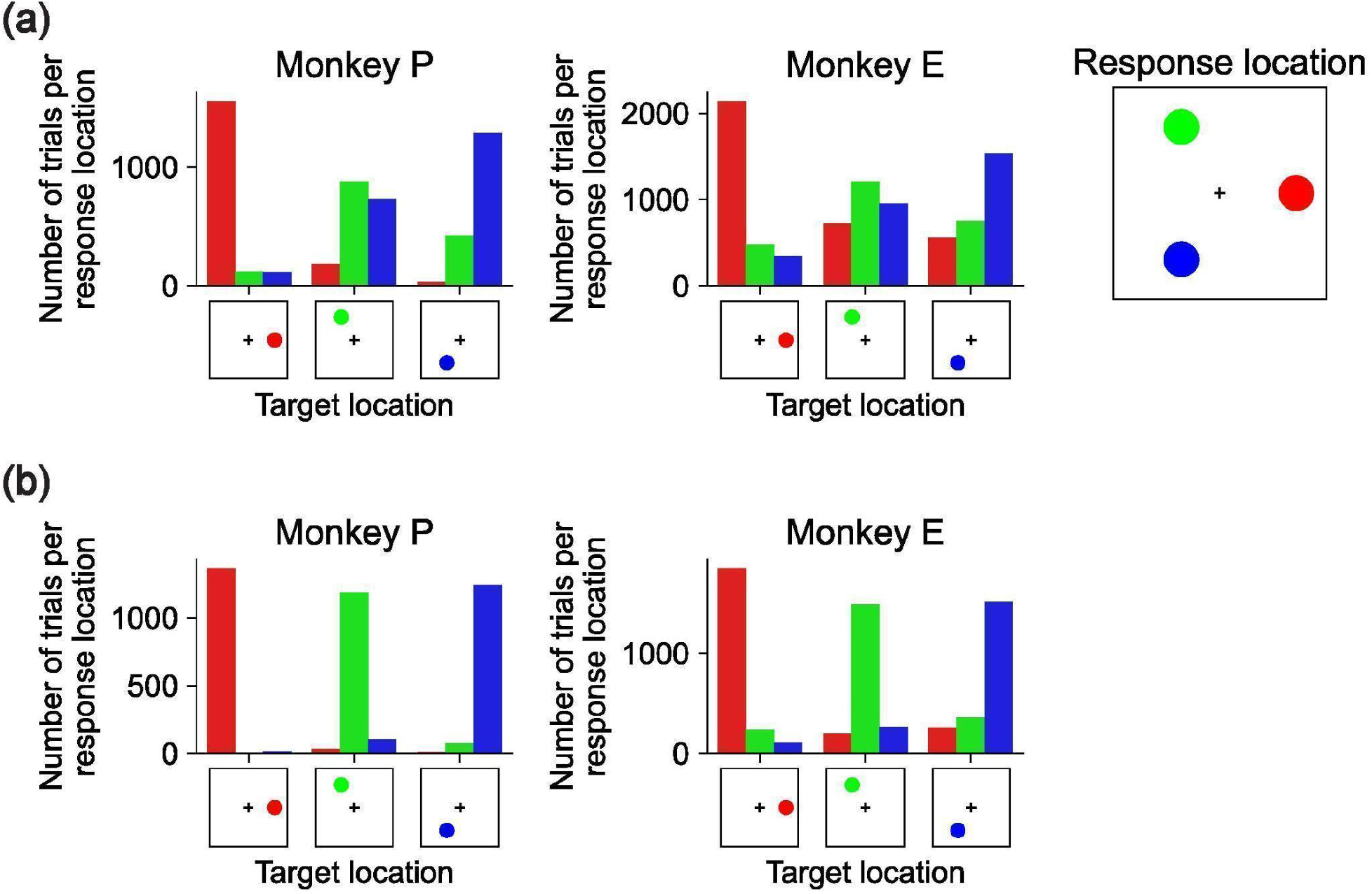
Triangle dataset error patterns. **(a)** 3-object trials for Monkey P (left) and Monkey E (middle). In each panel, the x-axis is the target location (the location of the cued object), the hue represents the location of the object that the monkey responds to according to the color scheme on the far right, and the y-axis is the number of trials across all sessions satisfying this pair of target and response locations. **(b)** Same as panel (a) but for 2-object trials.

**Extended Data Figure 2:**
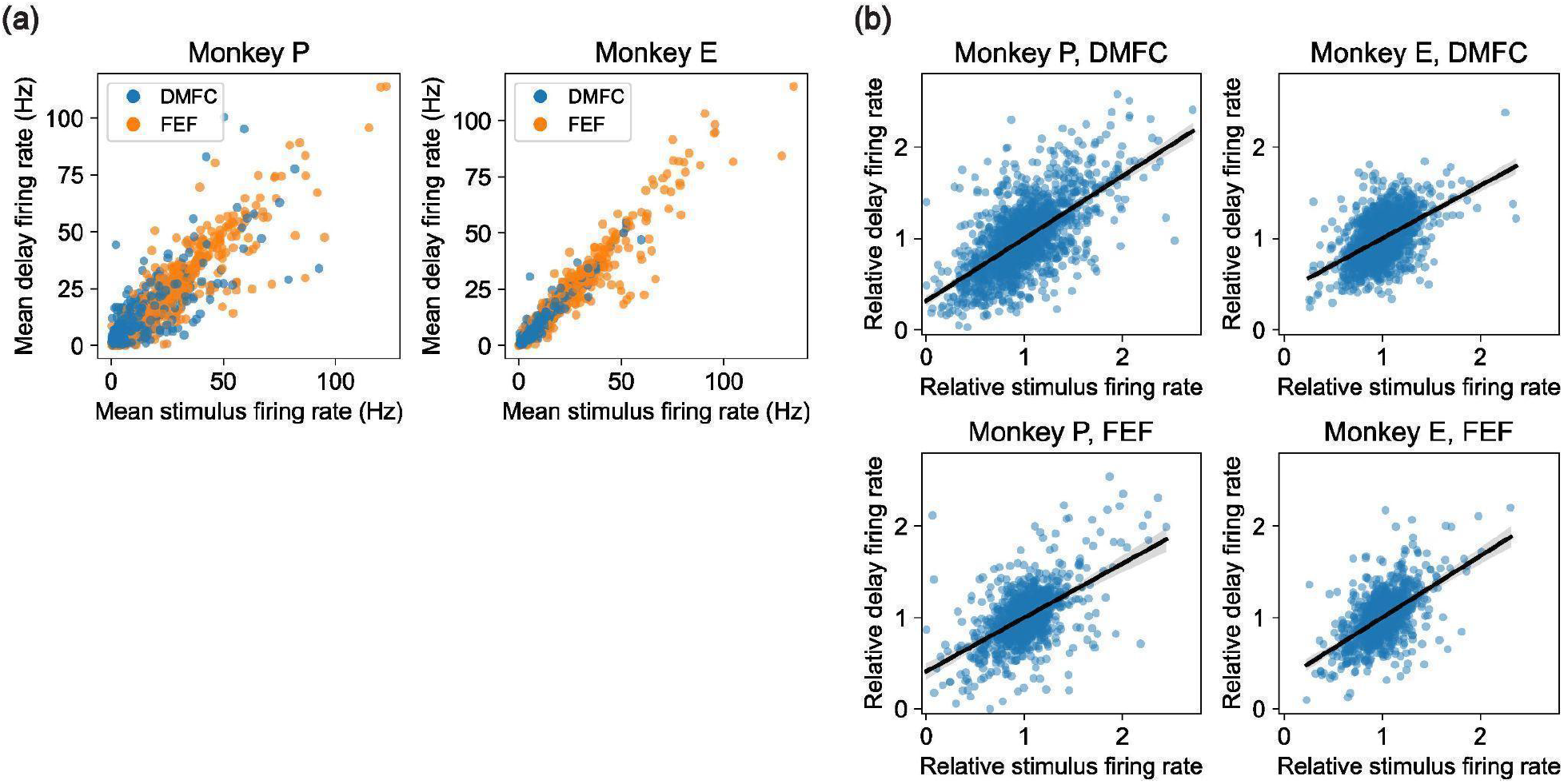
Firing rate statistics across stimulus and delay phases. **(a)** Stimulus vs delay phase firing rate per recorded unit for Monkey P (left) and Monkey E (right). Each point represents one neuron, colored by brain area (FEF in blue and DMFC in orange). X-axis is mean firing rate over the entire stimulus phase (0-1000ms after stimulus onset) and y-axis is mean firing rate over the entire delay phase (delay onset until cue onset). **(b)** Correlation in spatial selectivity across stimulus and delay phases for each recorded unit, for Monkey P (left column) and Monkey E (right column) in DMFC (top) and FEF (bottom). In each plot, every point corresponds to a pair [unit, location], for each recorded unit and each of the three locations in the triangle dataset. Given a unit and location, the x-axis value is the mean firing rate during the stimulus phase for that unit for 1-object trials where the object was at the given location, after scaling so that over all locations the unit has mean firing rate 1. This is a simple measure of how much the unit is selective to the given location over the other locations. The y-axis is analogous but for the delay phase.

**Extended Data Figure 3:**
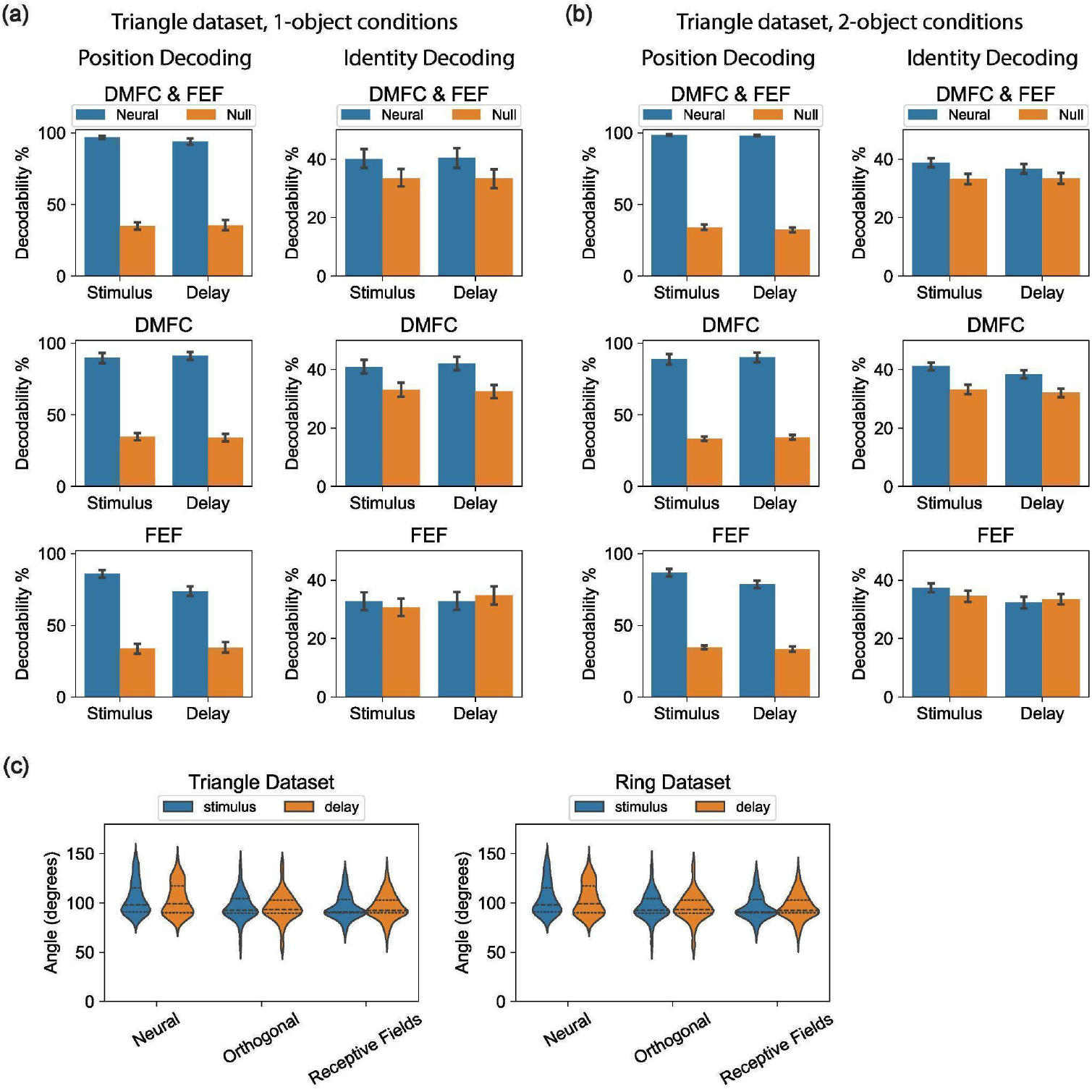
Decoding analyses. All results in this figure were generated from only the three recording sessions (from each of the Triangle dataset and Ring dataset) with the greatest number of units with significant task modulation (See Model-Fitting Methods), reflecting best-case results with the highest quality data we recorded. **(a)** Decoding Position (left) and Identity (right) for 1-object (top) conditions in the Triangle dataset using a linear decoder applied to single-trial firing rates from the stimulus and delay task phases (x-axis). Blue bars show results on neural data, and orange bars are results on neural data with randomly shuffled labels. Y-axis is cross-validated percent correct decoding, bootstrapped over ten 80%/20% train/test splits. The decoder was trained to output a one-hot 3-vector of the stimulus object’s location/identity. **(b)** Same as panel (a) except for 2-object conditions, and the decoder was trained to output a one-hot vector of the location/identity absent in the stimulus. Hence performance over 66% implies neural activity contains information about both objects. **(c)** Orthogonality of decoding axes for 2-object conditions in the Triangle dataset (top) and Ring dataset (bottom). The x-axis shows the angle between the neural coding axes of the two decoders in a 2-object decoder (see Methods), over all object locations and ten bootstraps. Results for the stimulus phase are shown in blue and the delay phase shown in orange. X-axis shows three datasets: The real Neural data, and two synthetic datasets with firing rate means and variances matched to the real data per unit. In the Orthogonal synthetic dataset, the two objects are forced to be encoded by two orthogonal subspaces in neural state space, and in the Receptive Fields synthetic dataset, both objects are encoded by the same set of neurons with spatial receptive fields (see Methods).

**Extended Data Figure 4:**
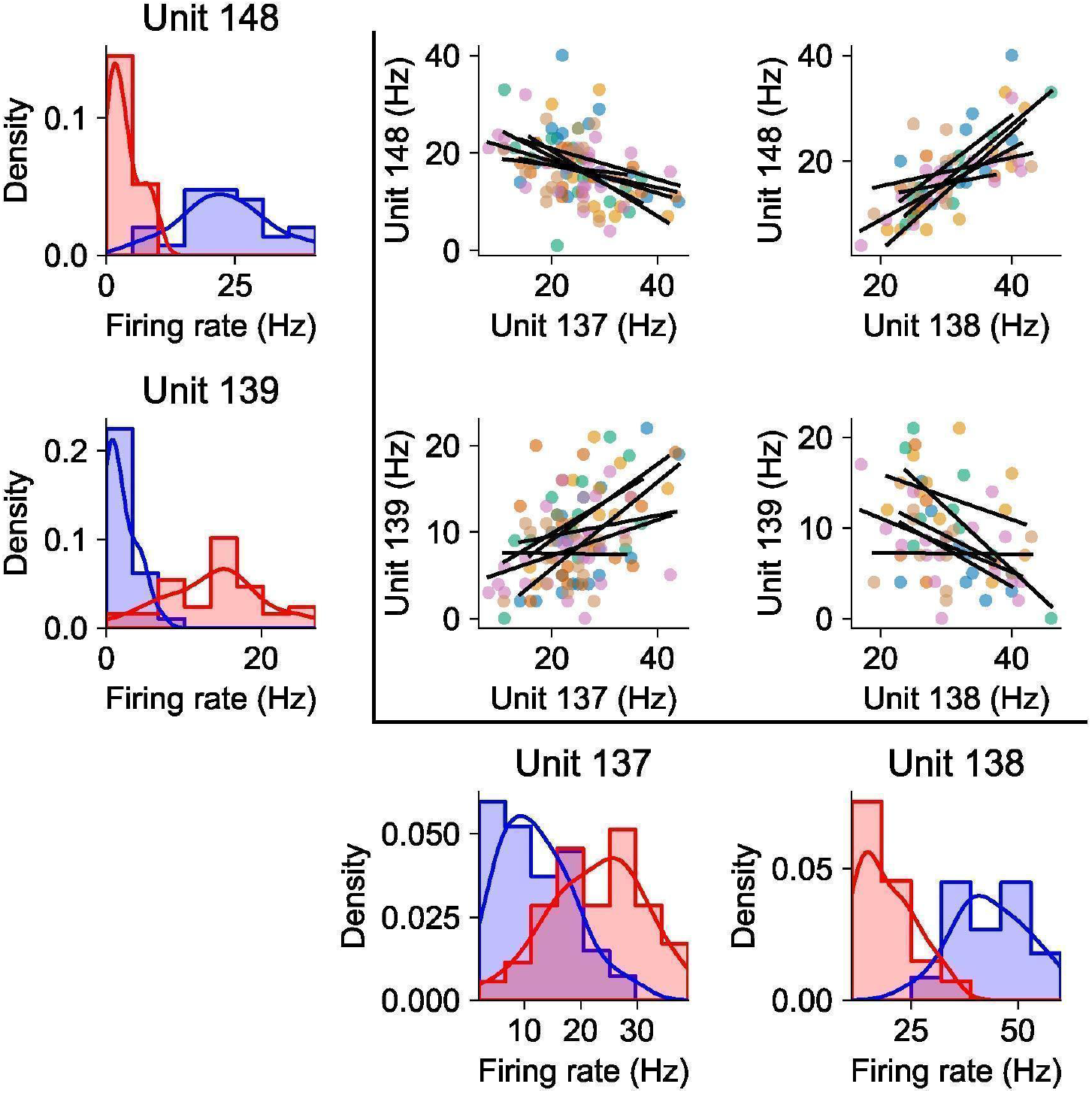
Example noise correlations conditionalized on identity. This figure shows the same data as Fig. 2c in the main text, except the scatterplots and regressions are conditionalized on stimulus identity. The colors in the scatterplots show each of the 6 identity combinations constituting different stimuli for the composite 2-object condition. Each black line shows the regression for one of these stimuli.

**Extended Data Figure 5:**
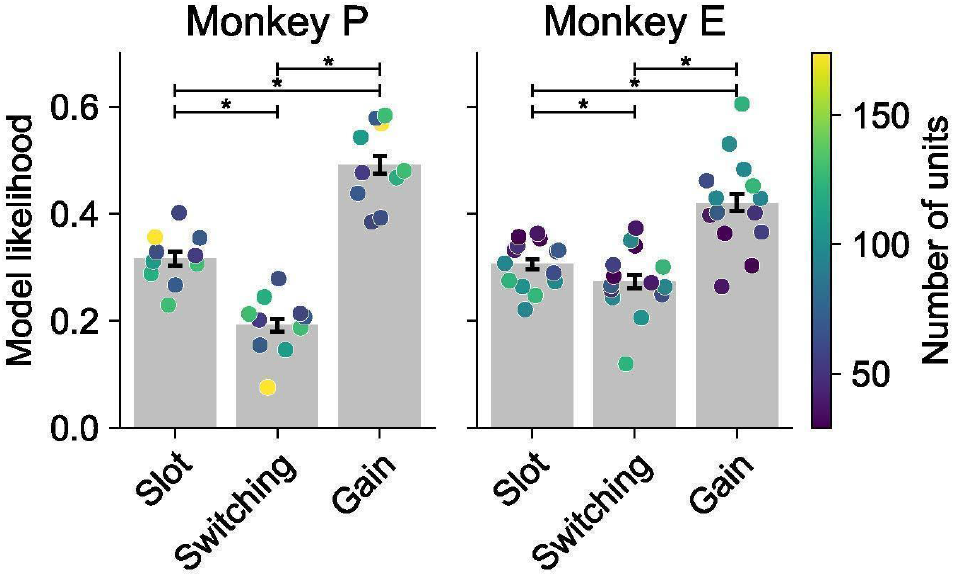
General Slot model results. Modeling results analogous to Fig. 4a, except using the General Slot model (see Methods) where each slot is allowed to be an arbitrary learned linear combination of neurons, instead of being a disjoint set of neurons.

**Extended Data Figure 6:**
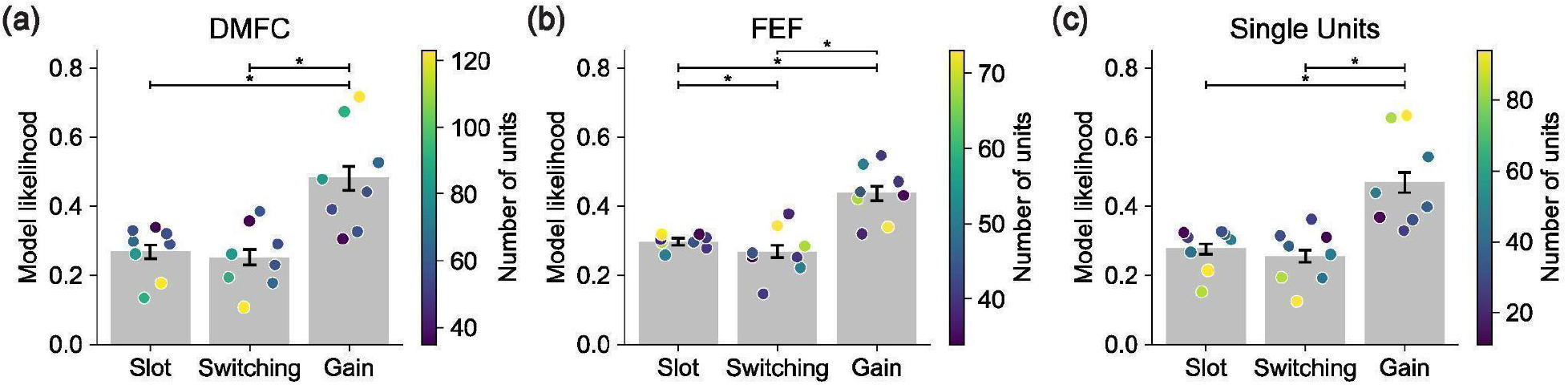
Control modeling results. **(a)** Modeling results analogous to Fig. 4a, except using only data from DMFC and only using 8 sessions (the four sessions with the most units from each monkey). **(b)** Analogous to panel (a) except for data from FEF. **(c)** Analogous to panel (a) except for data from only units labeled as single neurons.

**Extended Data Figure 7:**
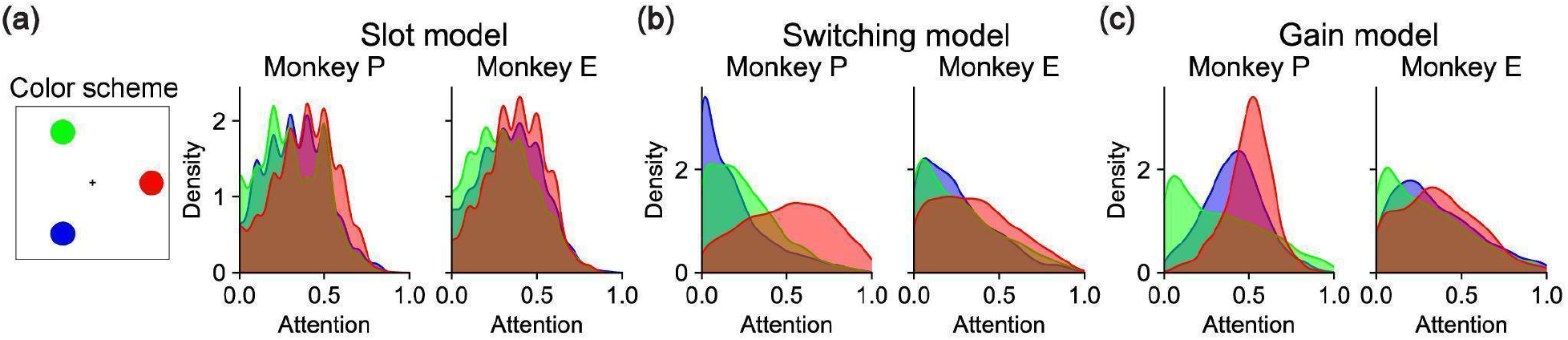
Distribution of attention. **(a)** *(left)* Color scheme used for other panels, assigning a color to each object location. *(right)* For each monkey, distribution of attention values from the Slot model on 3-object trials, conditionalized on object location as per the color scheme. Attention to an object from the Slot model on a given trial is zero if that object is not assigned to any slot and 1 if that object is assigned to a slot. **(b)** Analogous to panel (a) except for the Switching model, where attention to an object from the Switching model on a given trial is the mean fraction of timesteps for which that object is the object encoded by the model. **(c)** Analogous to panel (a) except for the Gain model, where attention to an object is the gain factor for that object.

**Extended Data Figure 8:**
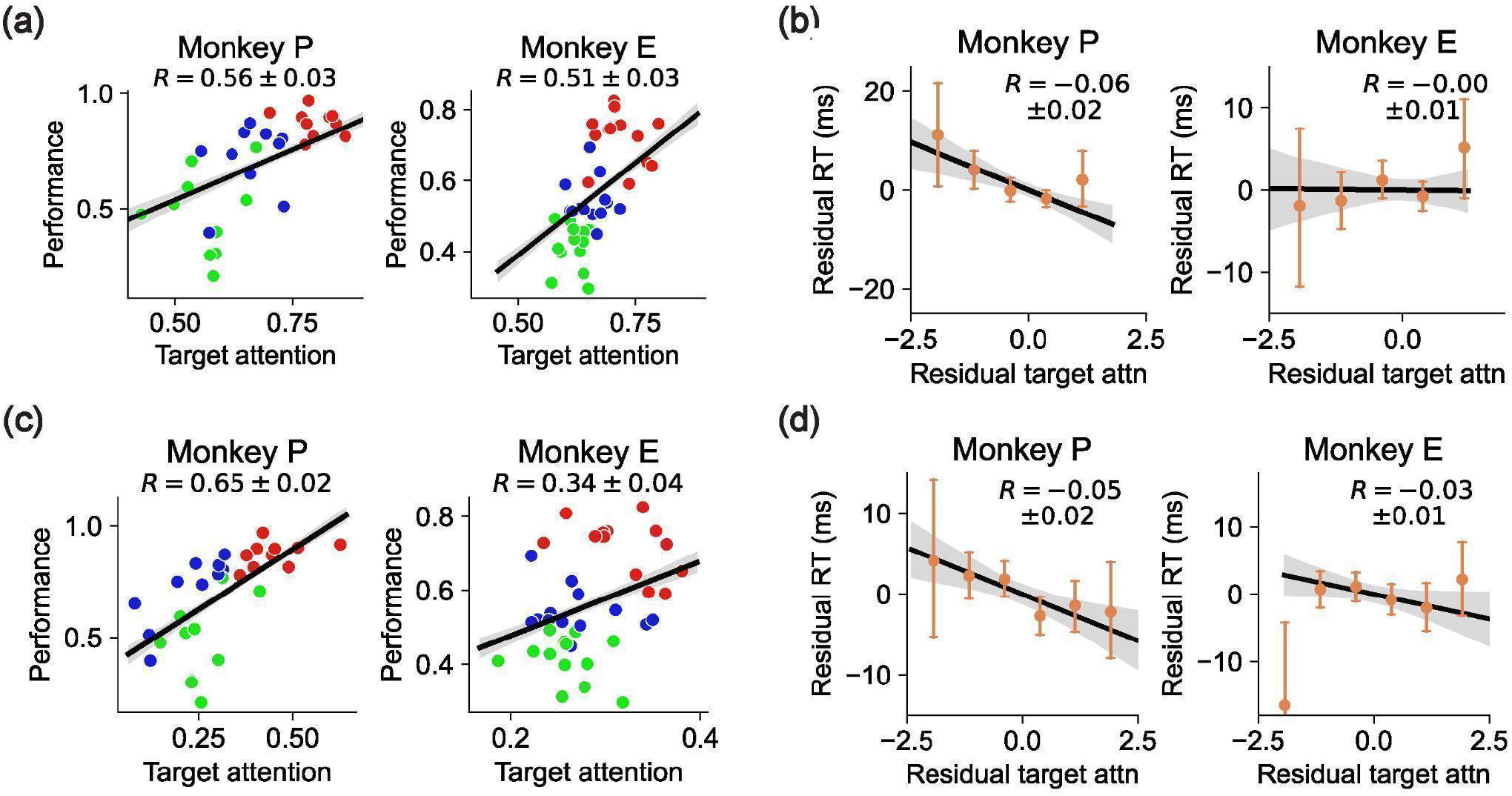
Behavioral predictions from Slot and Switching models. **(a)** Analogous to Fig. 4c except using attention from the Slot model on the y-axes. Attention to an object from the Slot model on a given trial is zero if that object is not assigned to any slot and 1 if that object is assigned to a slot. The regression is significantly positive for both monkeys (*R* = 0. 19±0. 05, *p* = 6⋅10^−4^ for Monkey P; *R* = 0. 36±0. 04, *p* < 10^−10^ for Monkey E). **(b)** Analogous to Fig. 4d except using normalized residual attention from the Slot model. The regression is not significant for either monkey (*R* = −0. 03±0. 017, *p* = 0. 034 for Monkey P; *R* = −0. 01±0. 014, *p* = 0. 40 for Monkey E). **(c)** Analogous to panel (a) except using normalized attention from the Switching model. Attention to an object from the Switching model on a given trial is the mean fraction of timesteps for which that object is the object encoded by the model. The regression is significantly positive for both monkeys ( *R* = 0. 65±0. 02, *p* < 10^−10^ for Monkey P; *R* = 0. 34±0. 04, *p* < 10^−10^ for Monkey E). **(d)** Analogous to panel (b) except for normalized residual attention from the Switching model. The regression is significantly negative for Monkey P but not significant for Monkey E (*R* = −0. 05±0. 016, *p* = 0. 002 for Monkey P; *R* = −0. 03±0. 014, *p* = 0. 063 for Monkey E).

**Extended Data Figure 9:**
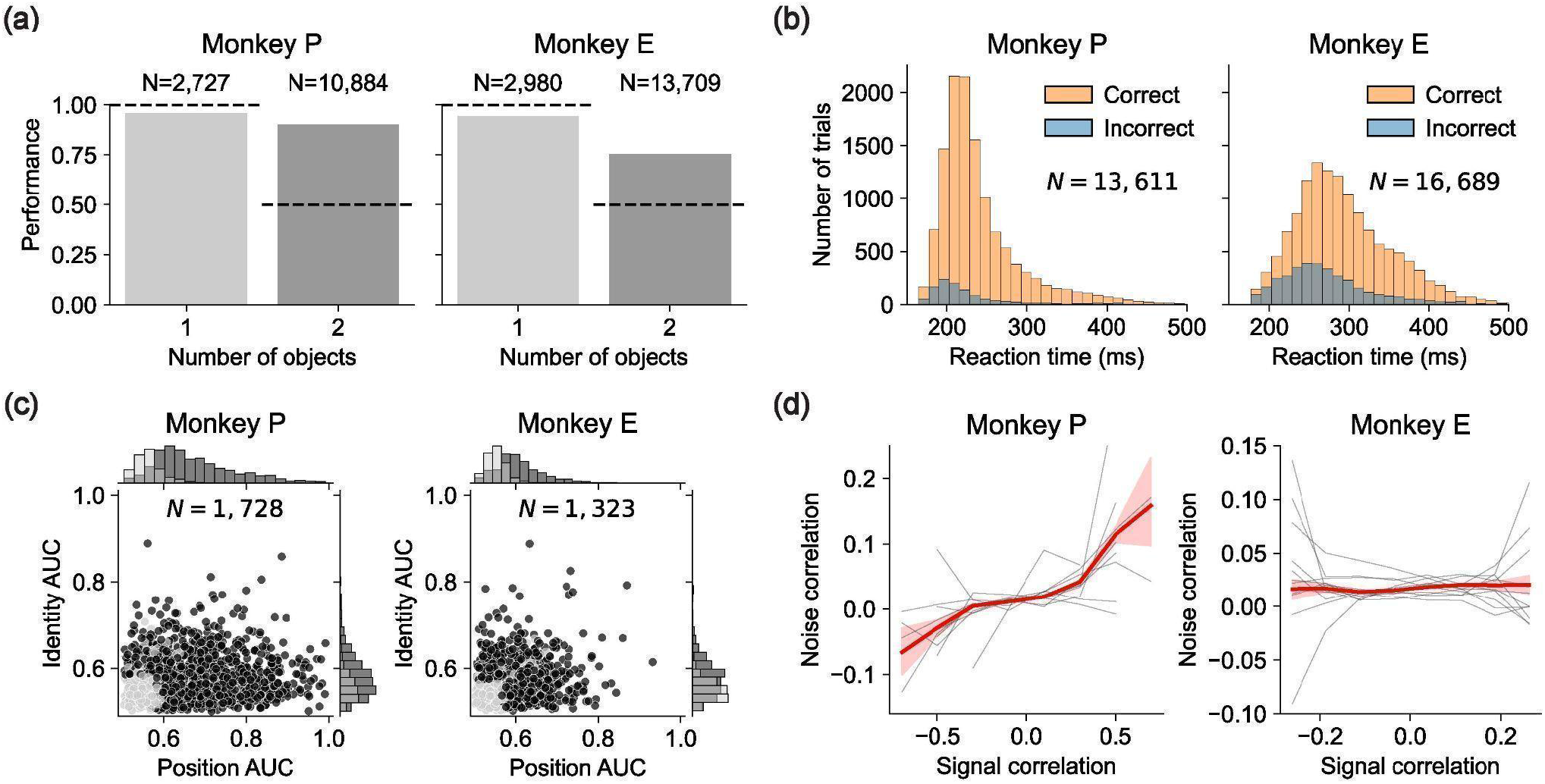
Ring dataset results per monkey. **(a)** Performance on the Ring dataset for Monkey P (left) and Monkey E (right), decomposing Fig. 5a per monkey. **(b)** Reaction times on the Ring dataset for Monkey P (left) and Monkey E (right), decomposing Fig. 5b per monkey. **(c)** Position- and identity-selectivity of every recorded unit in the Ring dataset for Monkey P (left) and Monkey E (right), decomposing Fig. 5d per monkey. **(d)** The relationship between noise and signal correlations for all pairs of simultaneously recorded neurons in the Ring dataset, for Monkey P (left) and Monkey E (right). This decomposes Fig. 5e per monkey.

**Extended Data Figure 10:**
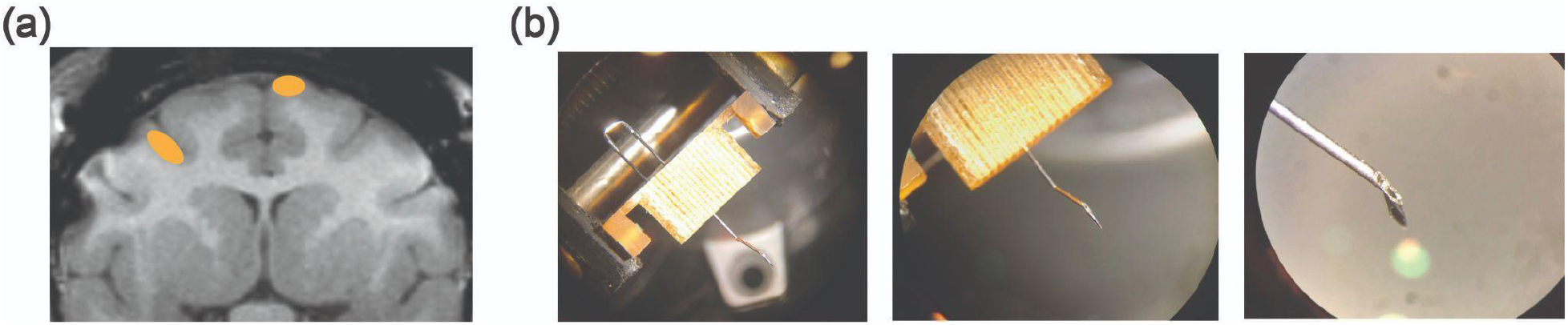
Recording targeting and guide tube system. **(a)** Coronal MRI slice of the brain of Monkey P. Orange ovals show approximate areas of DMFC (right) and FEF (left) that were recorded in this slice. This slice is near the most posterior part of DMFC we recorded, and near the most medial and anterior part of FEF we recorded. **(b)** Neuropixels recording guide tube, under three levels of magnification. The guide tube was made of a 23G syringe needle and hand-crafted with a Dremel and pliers. It passed through a grid and attached to a crimped J-shaped piece that then returned through the grid from the top to afford rotation stabilization. This was assembled and autoclaved before every session, then placed on the chamber for recording.

**Extended Data Figure 11:**
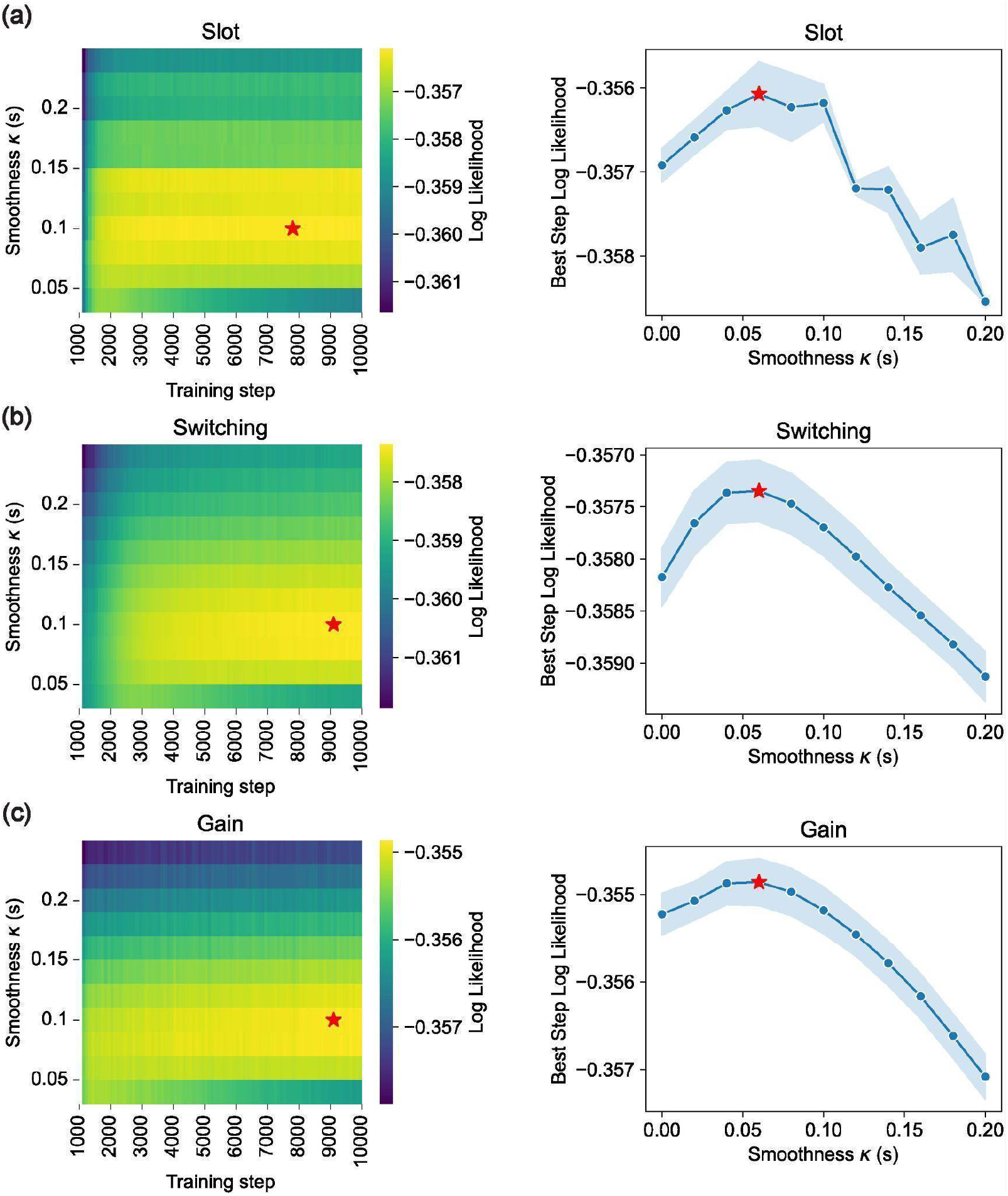
Model hyperparameter tuning. **(a)** (left) Heatmap of cross-validated log likelihood (hue) for the Slot model as a function of training step (x-axis) and smoothness parameter κ, for session 2022-06-01 of Monkey P. Red star shows point of maximum log likelihood. (right) Also for the Slot model on session 2022-06-01 of Monkey P, for each smoothness value (x-axis) we compute the training step with the highest log likelihood and show that log likelihood on the y-axis. Blue dots show mean over two random seeds, shaded region shows standard error of the mean, and red star shows maximum value. **(b)** Analogous to panel (a) except for the Switching model. **(c)** Analogous to panel (a) except for the Gain model.

## Notes

https://github.com/jazlab/multi_object_memory_2025

https://dandiarchive.org/dandiset/000620

https://osf.io/vyw49

### Competing Interest Statement

The authors have declared no competing interest.

### Summary of Updates

Minor correction to task description in Methods.

